# Preservation of Human Colonic Stem Cells Requires an ERK Dynamics Checkpoint Mediated by AKT

**DOI:** 10.64898/2026.04.02.715982

**Authors:** Lauren Riede, Alexander Borowiec, Saptarshi Mallick, Sohini Mallick, Jayati Chakrabarti, Curtis A. Thorne, Kelvin W Pond

## Abstract

Colonic stem cells reside in a microenvironment enriched in epidermal growth factor, which is essential for their survival and can activate both PI3K-AKT and MAPK-ERK pathways. This predicts co-activation of both pathways within the growth factor-high stem cell compartment at the base of crypts. However, in patient-derived human colonic organoids and normal human tissue, stem cells maintain robust AKT activity while suppressing ERK signaling despite active EGFR engagement. As stem cells differentiate, they activate pulsatile ERK signaling, which is essential for migration, survival, and maintenance of barrier function. We show that AKT-dependent phosphorylation of RAF-1 at serine 259 establishes a post-receptor checkpoint that maintains ERK temporal dynamics in stem cells. Acute activation of ERK in stem cells triggers rapid global differentiation. Disruption of the ERK checkpoint via mutation of serine 259 leads to sustained AKT and ERK co-activation in stem cells. Unlike ERK/AKT coactivation driven by apoptosis, co-activation in the stem cell compartment results in the emergence of a neoplastic, architecturally disorganized cell population dominating the cell fate profile. Incredibly, introducing brief ERK pulses through AKT inhibition or ERK activation triggers re-differentiation of neoplastic cells. Consistent with duration-dependent MAPK encoding principles, these data demonstrate that regardless of baseline signaling amplitude, ERK signaling dynamics are epistatic to total kinase signaling load in human colonic stem cells.

**SIGNIFICANCE:** Stem cells must balance self-renewal and differentiation while remaining responsive to continuous mitogenic stimulation to preserve tissue homeostasis. When self-renewal is impaired, wound healing and barrier integrity decline, whereas loss of proper differentiation drives tumorigenesis. Our findings demonstrate that this balance in the human colon is achieved through temporal control of kinase signaling rather than modulation of ligand availability. By establishing an AKT-dependent ERK dynamics checkpoint, colonic stem cells suppress differentiation-inducing ERK pulses while maintaining growth factor responsiveness. These results identify kinase dynamics as a fundamental determinant of epithelial homeostasis and suggest that subtle alterations in these dynamics may destabilize tissue organization during regeneration or chronic inflammation. Temporal encoding of kinase activity thus represents a central organizing principle in human stem cell biology.

## INTRODUCTION

The human colonic epithelium must continuously preserve the correct balance between two robust biological requirements: maintenance of long-lived adult colonic stem cells (CSCs) at the base of the crypt and continuous production of terminally differentiated epithelial cells above the stem cells which perform the essential functions of the colon^1, 2^. Here, we define epithelial homeostasis not simply as maintenance of proliferation or survival, but as stable preservation of lineage structure over time. CSCs do divide, but more slowly than their descendants in the transit-amplifying (TA) compartment, which undergo a burst of rapid proliferation before exiting the cell cycle and terminally differentiating into mature epithelial lineages^3^. Preserving this sustained pool of CSCs by restricting their entry into a TA state is therefore essential for lifelong renewal of this highly regenerative tissue and prevention of hyperplasia^4^.

A central unresolved question is how CSCs identity is preserved despite continuous exposure to noisy pro-differentiation factors. In the colonic epithelium, cell fates are distributed spatially along the crypt axis as are diffusible ligands secreted from the underlying stroma that promote stemness, ^5, 6^. Epidermal growth factor (EGF) is one such ligand, essential for epithelial survival, which triggers rapid EGFR activation followed by co-activation of both the PI3K-AKT and MAPK-ERK signaling pathways in colonic epithelial cells^7, 8^. However, here we show that *ex vivo* PDCOs and *in vivo* human colonic tissue do not uniformly co-activate these pathways in a gradient originating from the crypt base as one might expect. Instead, AKT and ERK signaling are spatially partitioned into largely mutually exclusive regions, with ERK activity being strongly excluded from LGR5+/MYC+/SOX9+ stem-cell compartments despite abundant EGF and EGFR. Thus, shared dependence on EGFR does not translate into shared downstream signaling output, arguing that colonic epithelial cells actively partition receptor output according to cell state rather than passively mirroring a diffusible ligand gradient. We describe **signaling insulation** as the ability of CSCs to maintain selective downstream pathway activation despite shared extracellular input. Well established examples of this include receptor partitioning during GPCR desensitization^13^, sequestration of RTK monomers^9, 10^, competing Erb-family heterodimers^11^, and Wnt receptor regulation by R-spondin, LGR5, and RNF43, which are required to maintain CSCs^12,13^.

Here, we use the term **ERK checkpoint** to describe a new mechanistic arm of signaling insulation in CSCs: AKT inhibits RAF-1 via phosphorylation at serine 259, which restrains downstream ERK activation in stem cells and preserves feedback-limited ERK dynamics. We propose that this checkpoint enables ERK to function as a transient, pulse-responsive signal rather than a sustained mitogenic program reliant on ligand concentration. When intact, the checkpoint permits brief ERK activation that can drive the coordinated behaviors required for epithelial repair, including transient hyperproliferation, collective movement, and subsequent differentiation. When this buffering mechanism fails, mitogenic input is converted into aberrant sustained AKT/ERK co-activation associated with proliferation and migration, dysplastic patterning, and what others have shown is pro-neoplastic kinase signaling^14^. Thus, the ERK checkpoint is required to preserve the ability of the epithelium to use ERK dynamically without collapsing into a pathological signaling state.

This distinction is important because ERK signaling plays essential physiological roles in the colon and is not inherently detrimental. We and others have shown it is required for migration, wound healing, and maintenance of gut barrier function^8, 15–19^, meaning that homeostasis cannot be maintained through global suppression of ERK signaling. Instead, the subset of stem cells must selectively buffer pro-differentiation inputs to preserve their identity and prevent inappropriate fate transitions. Failure of this buffering system would be expected to destabilize the tissue in two opposite but pathological directions: excessive differentiation that depletes the stem-cell pool and impairs repair ability, or chronic hyperactivation that promotes dysplastic remodeling and may increase susceptibility to neoplasia^20–22^.

Here, using patient-derived human colonic organoids and normal human tissue, we identify signaling insulation as a core organizing principle of epithelial homeostasis. We show that although EGF/EGFR signaling is required broadly across the epithelium during homeostasis, activation of downstream effectors are partitioned into two distinct pools: 1. Activation of AKT and suppression of ERK uniformly in stem cells and 2. Sporadic co-activation of AKT and ERK in clusters of differentiated cells. We establish that AKT-dependent phosphorylation of RAF-1 at S259 drives an ERK checkpoint in stem cells to preserve feedback-limited pulsatile ERK dynamics. Sustained pathway hyperactivation without pulsatile dynamics, when the ERK checkpoint is abolished, expands a neoplastic population of AKT/ERK co-activated cells associated with aberrant patterning and ERK dynamics frequently observed in colorectal cancers^14^. Together, these findings show that colonic homeostasis is governed not simply by pathway strength, but by how EGFR outputs are partitioned across cells and encoded over time. Importantly, acute AKT inhibition or ERK activation re-introduces a transient ERK pulse that is sufficient to drive differentiation even in the setting of oncogenic driven ERK hyperactivation, indicating that in this setting, ERK signaling dynamics are epistatic to total kinase signaling load.

## RESULTS

### Colonic Stem Cells Suppress ERK Despite Ample EGF/EGFR Availability

Patient-derived colonic organoids (PDCOs) maintain unique kinase signaling and high-fidelity cell type heterogeneity even when exposed to the uniform and excess mitogens such as EGF. PDCOs can also repeatedly self-organize into a patterned architecture of heterogenous cell types that mimic *in vivo* human colon physiology in both 3D and 2D organoid culture systems (Fig. 1A). Self-organized organoids are classified as homeostatic when cells fully polarize (Fig. S1A) and fate markers stabilize into three temporally stable and quantifiable niches (Fig. S1B), despite the high turnover of the colonic epithelium.

1. Stem Cells-OLFM4⁺/Sox9⁺ dense clusters of cells (∼5-10%) (Fig. 1B, S2A).
2. TA Progenitors-Ki67⁺/EdU⁺ transit-amplifying cells (∼5-10%) (Fig. 1C, S2B).
3. Differentiated Cells-CK20⁺ differentiated colonocytes (∼80-90%) (Fig. 1D).

**Figure 1:**
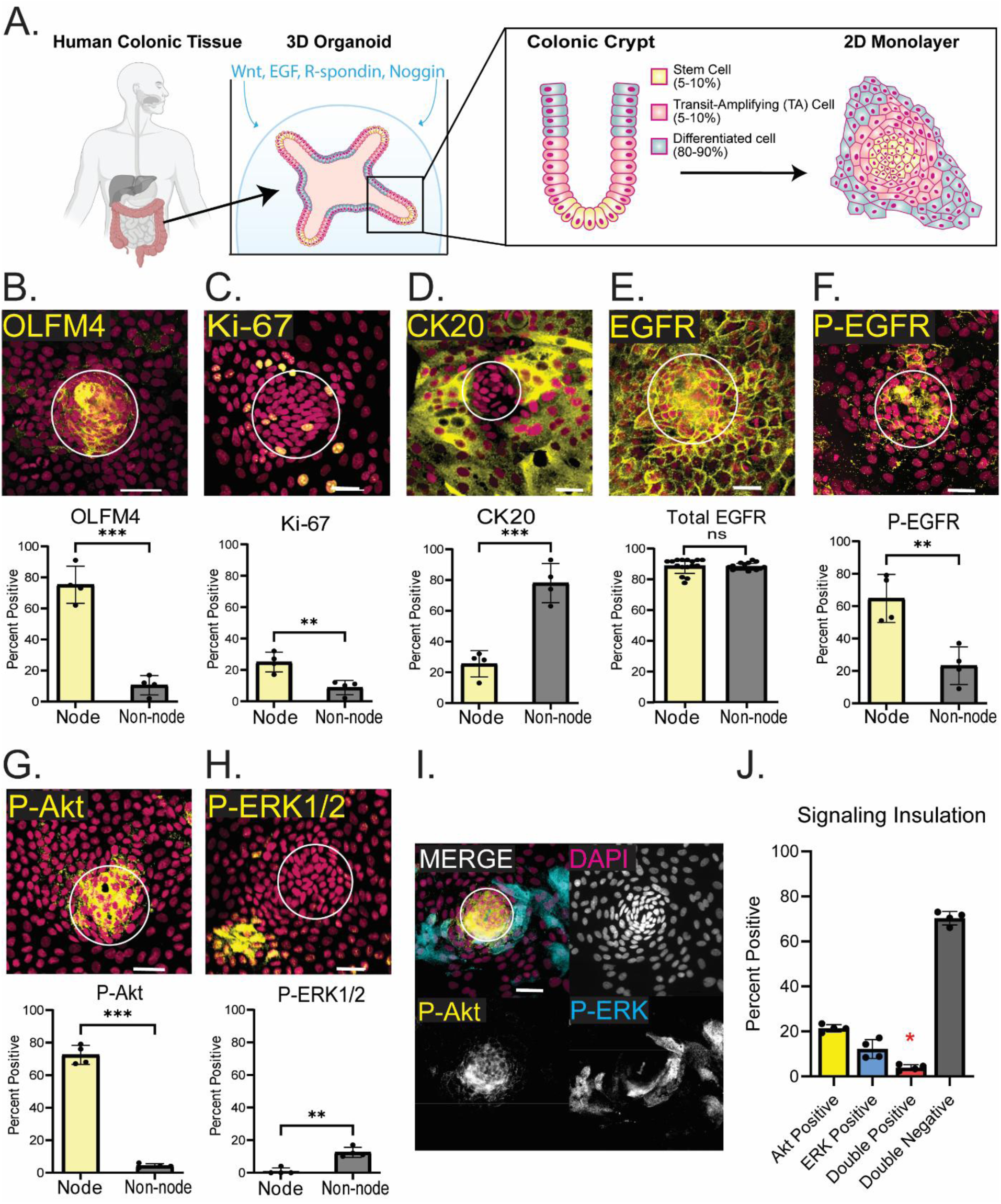
Colonic Stem Cells Suppress ERK Despite Ample EGF/EGFR Availability. **(A)** Model depicting organoid monolayer preparation. Patient colonic tissue is digested, single celled, and plated in a bubble of Matrigel. Organoids are supplemented with media containing surplus of Wnt, EGF, R-spondin, and Noggin. Once organoids are fully grown in 3D conditions, cells are disassociated and plated on a thin layer of Matrigel. Within 5-7 days, cells grow into a monolayer with distinct stem cell (5-10% of cells), TA cell (5-10% of cells) and differentiated cell (80-90% of cells) niches. Stem cell niches, called nodes, are dense, regularly spaced niches made up of OLFM4/Sox9/LGR5/MYC positive stem cells. **(B)** Representative image of stem cell niche (node) characterized by OLFM4+ cells with quantification. OLFM4 measured through immunofluorescent staining, population makes up around 10% of monolayers, although almost all cells in node are OLFM4+. Cells were portioned into node or non-node and percent of OLFM4+ cells out of total node or non-node cells was used for quantification. **(C)** Representative image of transit-Amplifying (TA) cells characterized by Ki-67 immunofluorescent staining. TA cells surround the OLFM4+ stem cells and make up around 10% of the monolayer. **(D)** Differentiated cells shown by CK20 immunostaining makes up the surrounding monolayer, around 80% of total cells. **(E)** Total EGFR representative image and quantification of almost all cells in monolayer expressing EGFR uniformly. **(F)** P-EGFR-Y1068 representative image and quantification preferentially activated in nodes. **(G)** Representative image and quantification of P-AKT-S473 exclusively activated within nodes and low in differentiated compartment. **(H)** P-ERK1/2 representative image and quantification. Activation excluded from nodes and on in a small percentage of differentiated cells at any given time (10-20%). **(I)** Co-stain representative image of P-AKT and P-ERK mutual exclusivity. **(J)** Quantification of AKT+, ERK+, Double Positive and Double Negative cells within monolayer. Red asterisks marking around 5% total cells that are double positive, uninsulated cells that coactivate AKT and ERK. Three to four biological replicates were performed. Data shown is from analysis of 4 technical replicates with at least 150 total cells quantified per replicate. Data are represented as mean ± SEM. All scale bars are 100um, significance calculated with Welch’s t-test, ** P ≤ 0.01, *** P ≤ 0.001. See also Figure S1 and S2.

EGF activated EGFR is essential for survival of epithelial cells in the gut and a strong activator of both AKT and ERK kinase pathways. Despite equal total receptor levels (Fig. 1E), PDCOs grown in 100ng/mL EGF, display sporadic co-activation of AKT, ERK1/2, and EGFR in a minority of differentiated cell clusters (Fig. 1F, S2C-D) consistent to what has been shown previously using clonal cell lines^8^. This signaling pattern was in contrast to the dense stem cell compartments, which showed uniformly high AKT levels and was absent of ERK activation (Fig. 1G-I). Treatment with 500nM Gefitinib, a potent EGFR inhibitor, suppressed both AKT in the stem cells and ERK/AKT in the differentiated compartment by 10-20 fold after 1hr incubation (Fig. S2E), indicating that both unique kinase cascades are reliant on EGF/EGFR stimulation. In order to define how these bifurcated pathways maintain signaling insulation via an ERK checkpoint, we quantified the ratio of cells co-activating AKT/ERK, which are present at <5% in normal homeostatic cultures (Fig. 1J-red asterisk).

We next examined AKT and ERK activation in normal human colonic tissue sections taken during colonoscopy to ensure *in vivo* CSCs were signaling similarly as our *ex vivo* PDCO model. Despite a gradient of cell fates in the colonic crypts (Fig. S3A-C), pAKT localized exclusively to the crypt base and pERK was restricted to the differentiated compartments and regularly spaced across the epithelium in wave-like patterns (Fig. 2B). IHC images and crypt-axis quantification support spatial segregation of PDCOs and partitioning of ERK signaling exquisitely mimics what is seen in normal human tissue *in vivo* (Fig. 2B-D). The distance between crypts (Fig. 2c’), between ERK activity clusters (Fig. 2c’’), and between a crypt and its nearest ERK-active cluster (Fig. 2c’’’) was remarkably conserved (Fig. 2D). The size of ERK waves within normal colonic tissue is roughly one half what is measured through live-cell imaging of ERK dynamics in organoid monolayers, due to the loss of temporal information in the IHC staining (Fig. 2D) These data show that the 2D PDCO platform is high fidelity and phenocopies the patterning, tissue level architecture, cell shape differences, and most importantly the signaling insulation observed in normal human colonic epithelium *in vivo*.

**Figure 2:**
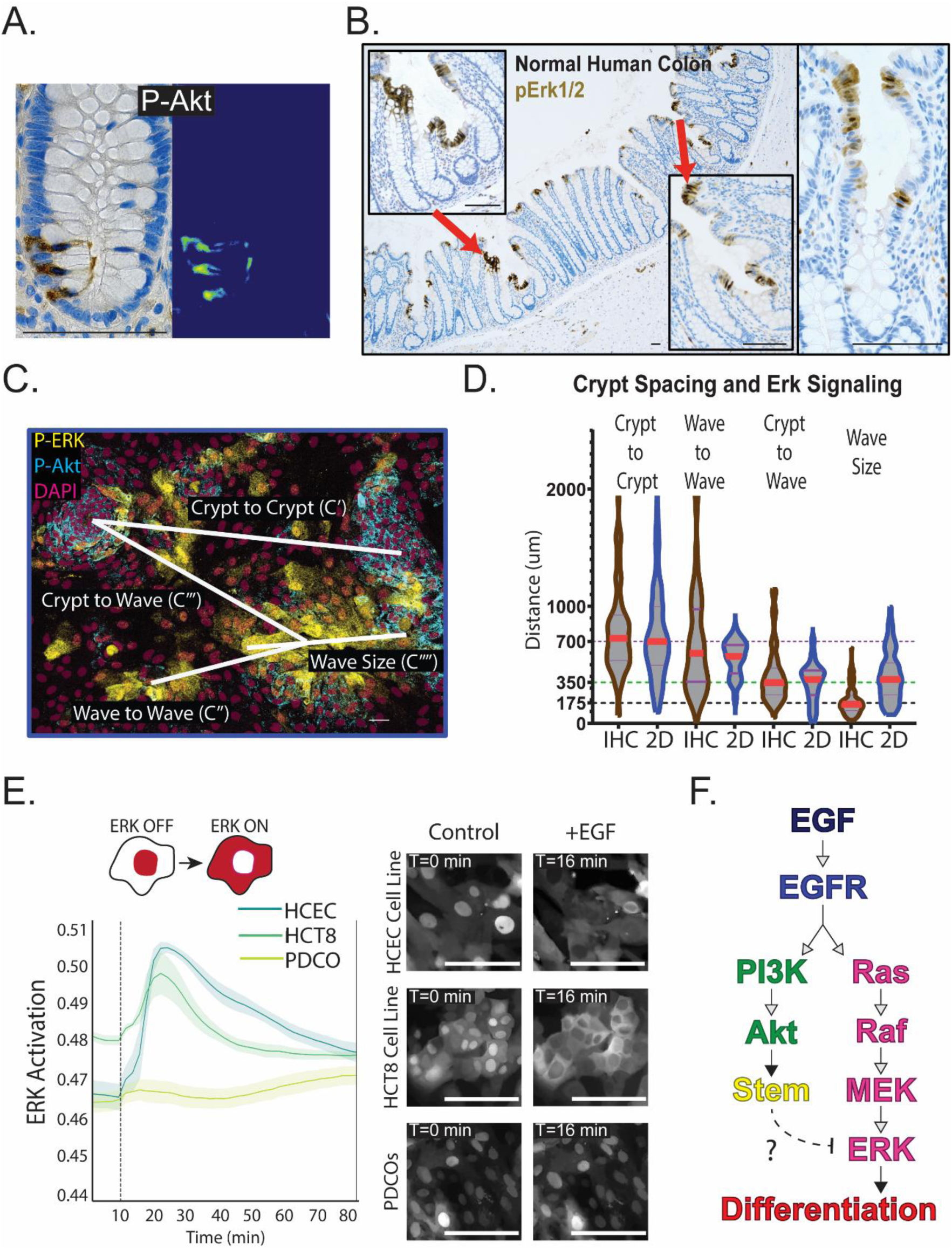
Colonic Stem Cells Suppress ERK Despite Ample EGF/EGFR Availability. (**A**) Representative image of normal human colonic crypt immunohistochemistry of P-AKT within the stem cell compartment specifically. **(B)** Immunohistochemistry staining of P-ERK1/2 in normal human colonic tissue with zoomed in regions showing activation exclusively in the top, differentiated, compartment of crypts. **(C)** Representative image of P-AKT and P-ERK immunostaining of PDCO monolayer showing spacing of crypt to crypts (C’), crypts to waves (C’’), and wave to waves (C’’’), and wave size (C’’’’). **(D)** Quantification of IHC staining (brown violin plots) and organoid monolayer staining (blue violin plots) of P-ERK. Individual crypts IHC images and PDCO monolayers were measured for crypt to crypt, ERK wave to ERK wave, crypt to ERK wave, and total wave size. **(E)** 100ng/mL EGF pulsed into human colonic cell lines HCEC and HCT8 and organoid monolayers expressing ERK-KTR biosensor and H2B-iRFP nuclear marker. Data shown as mean activation, calculated through nuclear/cytoplasmic ratio of biosensor intensity with ribbons depicting 95% confidence intervals. 3 biological replicates were performed, analysis completed on 4 technical replicates with at least 150 cells quantified per condition. **(F)** Model showing EGF activation of EGFR, which canonically activates both PI3K-AKT pathway and MAPK-ERK pathway. In organoids, MAPK-ERK pathway is suppressed within organoid stem cells despite EGFR activation. All scale bars are 100µm. Violon plot shows all measurements with bars representing mean ± SEM. See also Figure S3.

PDCO stem cells display a unique response to EGF when compared to epithelial cell lines, which rapidly co-activate ERK and AKT signaling. To determine if continuous exposure to EGF within the media led to ERK1/2 shutdown through negative feedback, we compared PDCOs and 2 immortalized colonic cell lines, normal human colonic epithelial cells (HCEC)^23^ and colonic adenocarcinoma cells (HCT-8), in their response to starvation followed by acute dosing of 100ng/mL EGF. ERK dynamics were evaluated for all three models using the well-established ERK-kinase translocation reporter (ERK-KTR)^24^. ERK activity is evaluated by the localization of the fluorescent reporter, which translocates from the nucleus to the cytoplasm when ERK1/2 is activated (Fig. 2E, top). ERK dynamics are quantified through single cell tracking and measurement of the cytoplasmic/nuclear ratio of reporter intensity. Normal HCEC cell lines and transformed HCT8 cell lines exhibited a robust ERK response to EGF within minutes while PDCOs showed almost no response to EGF stimulation (Fig. 2E, bottom). Incubation with alternative EGFR ligands was unable to activate ERK (Fig. S3D, E), indicating that EGF is the dominant ligand driving EGFR-dependent activation of ERK and AKT. These data suggest that unlike differentiated cells or cell lines, CSCs actively suppress ERK while maintaining EGFR/AKT activity, suggesting the presence of a post-receptor insulation mechanism to prevent aberrant differentiation (Fig. 2F).

### A Single ERK Pulse Triggers Robust Differentiation Program Independent of Baseline Signaling

To determine if post-receptor insulation of ERK is required to maintain stem cells, PDCOs were plated into a monolayer and grown until fully organized and patterned, around 6 days. Monolayers were then evaluated after one hour treatment with pharmacological perturbation for short-term signaling changes and 72 hours after treatment to evaluate long-term cell fate changes (Fig. 3A). To induce acute EGFR independent activation of ERK1/2, PDCOs were treated with 100nM of a PKC activator, Phorbol 12-myristate 13-acetate (PMA/TPA) for one hour then washed out. PMA-treated cells within nodes and outside of nodes both activated ERK and cells within nodes lost P-AKT activity completely (Fig. 3B-C, S4A-B). Despite hyperactivation of ERK after PMA treatment, no change was observed in signaling insulation (Fig. 3D), indicating that the ERK checkpoint remained intact following treatment. Changes in ERK dynamics in <150 single cells over a 24-hour period showed a strong induction of ERK activation immediately following addition of PMA (>4 hours), followed by a return to baseline ERK levels for the remainder of the 24 hours (Fig. 3E). Single cell quantification confirms the global 4-hour pulse of high ERK activity induced by PMA was ubiquitous across cells and did not result in any cell death after 24hrs (Fig. 3F, S4C-D). Coordinated oscillations of ERK in control cells is due to the regular timing of ERK waves driven by normal turnover of differentiated cells as shown previously^8, 19^. To quantify the extent of overall dynamics a cell experiences over time regardless of absolute signaling level (Kinase Load Score-KLS), we calculated the standard deviation of signaling for each cell, which we termed the Kinase Dynamics Score (KDS). In contrast to the insulation score, KDS increased dramatically in PMA treated cells (Fig. 3G). To measure the effect of this short pulse of ERK on the eventual cell fate, cell fates were quantified by IF staining of OLFM4+(Stem), Ki67+(TA), and OLFM4-Ki67-(Differentiated) (Fig.3H). Strikingly, three days after the pulse of ERK activity was initiated, the tissue exhibited a 2-fold reduction of stem cells and a 3-fold reduction in TA cells (Fig. 3I). Signaling insulation and KDS scoring indicates that ERK hyperactivation does not in itself de-insulate cells, and that KDS score could be an accurate predictor of long-term cell fate, as high KDS correlates here with CSC differentiation. These data show that a short-term pulses of ERK can function as an acute differentiation trigger in CSCs (Fig 3J).

**Figure 3:**
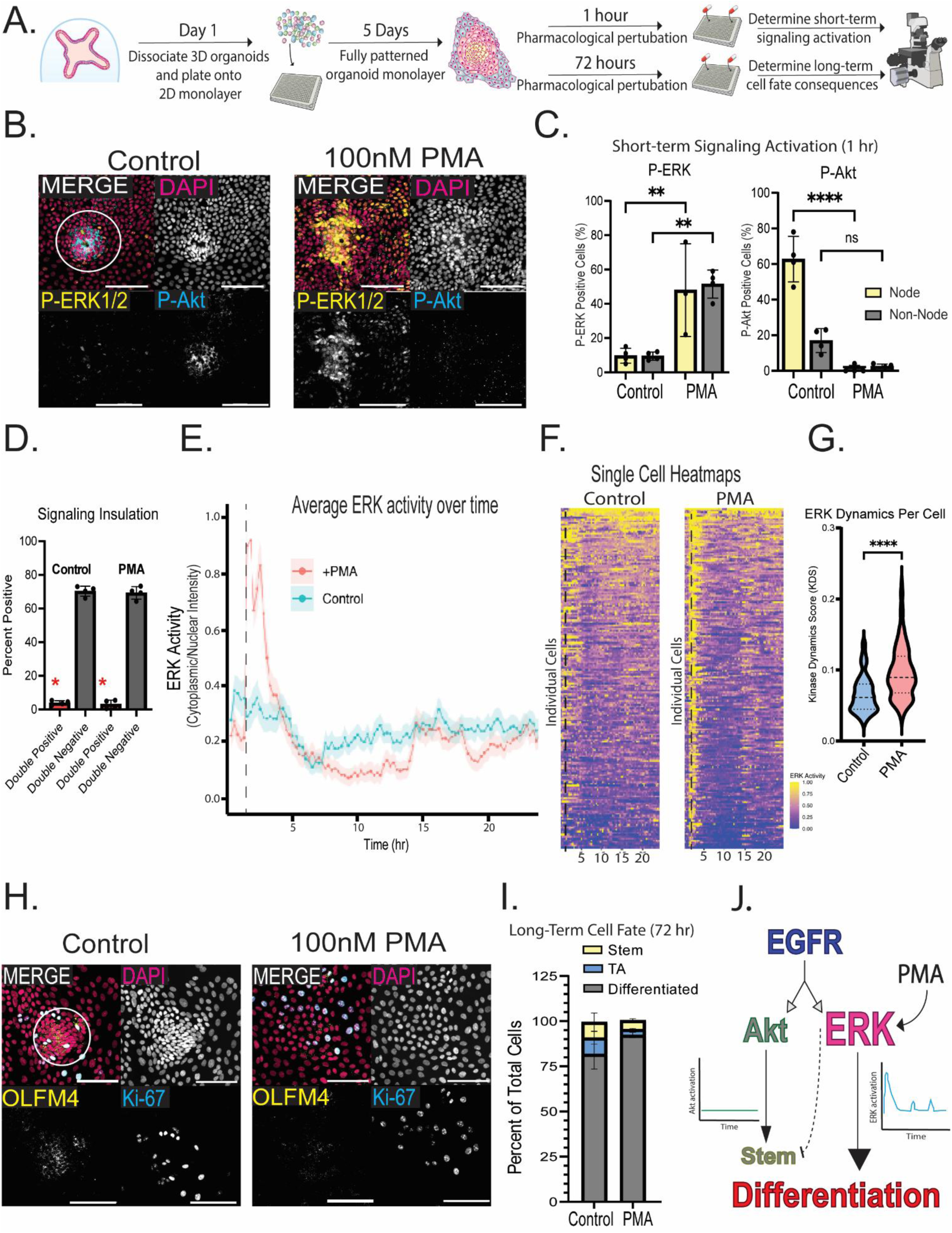
A Single ERK Pulse Triggers Robust Differentiation Program Independent of Baseline Signaling. **(A)** Experiment schematic of how pharmacological perturbations are tested. Fully grown 3D organoids are dissociated into single cells and plated into a 384-well plate covered in a thin layer of Matrigel. Organoids are grown for at least 5 days, or until cells make a full monolayer with proper node patterning. Wells are pharmacologically treated for 1 hour to look at short-term signaling activation changes (P-ERK and P-AKT) or 72 hours to assess long-term cell fate consequences to allow proper time for cell differentiation. **(B)** Immunofluorescence staining of P-ERK1/2 (yellow) and P-AKT (blue) in control and 100nM PMA treated cells. **(C)** Quantification of percentage of cells in node and outside of nodes with active ERK and AKT. **(D)** Quantification of ERK/AKT double positive cells and double negative cells. Red asterisks indicate less than 5% of cells activate both pathways, signaling insulation which is maintained in the presence of PMA. **(E)** Quantification of ERK-KTR biosensor in organoids treated with PMA and control. Organoids were treated with 100nM PMA 1 hour after starting the movie. Graph shows average ERK activity by cytoplasmic/nuclear ration of KTR intensity with ribbon showing 95% CI. **(F)** Heatmaps showing ERK activity of all cells tracked, with each cell being a horizontal line on the graph. Yellow shows high ERK activity and blue being low activity. **(G)** Kinase dynamics score (KDS) violin plot calculated as the standard deviation of ERK activity within each cell over the first 5 hours of the movie. **(H)** Immunofluorescence staining of cell fate markers OLFM4 (yellow, stem cells) and Ki67 (blue, TA cells) in control and 100nM PMA treated cells. **(I)** Quantification of percent of total cells that are stem or TA cells showing PMA causes a two-fold loss of stem cells. **(J)** Model showing PMA causes a pulse of ERK activity, leading to loss of AKT, loss of stem cells, and global differentiation of the monolayer. Three to four biological replicates were performed. Data shown is from analysis of 4 technical replicates with at least 150 total cells quantified per replicate. Data are represented as mean ± SEM. All scale bars are 100µm, significance calculated with four-way ANOVA, ** P ≤ 0.01, **** P ≤ 0.0001. See also.

### AKT-Dependent RAF-1-S259 Phosphorylation Suppresses ERK Signaling in Stem Cell Niches

MAPK signaling in epithelial cell lines have shown that phosphorylation of RAF-1 at serine 259 by AKT promotes binding of 14-3-3 to trap RAF1 in an inactive conformation^25, 26^. To investigate this as a potential mechanism controlling the insulation of ERK within stem cells, we quantified RAF1-S259 phosphorylation levels in normal human colonic tissue and 2D organoid monolayers. Both *in vivo* and *ex vivo*, phosphorylation of RAF1 at inhibitory site S259 was significantly higher in stem cell niches than in differentiated compartments (Fig. 4A-D). To confirm that AKT is driving phosphorylation of RAF-1, we treated PDCOs for one-hour with the AKT inhibitor, MK-2206. After treatment for 1 hour, RAF-1-S259 was reduced by 3-fold specifically within stem cell niches (Fig. 4E-F, S5A), suggesting that AKT directly enforces suppression of RAF-1 in stem cell niches to inhibit ERK signaling (Fig. 4G).

**Figure 4:**
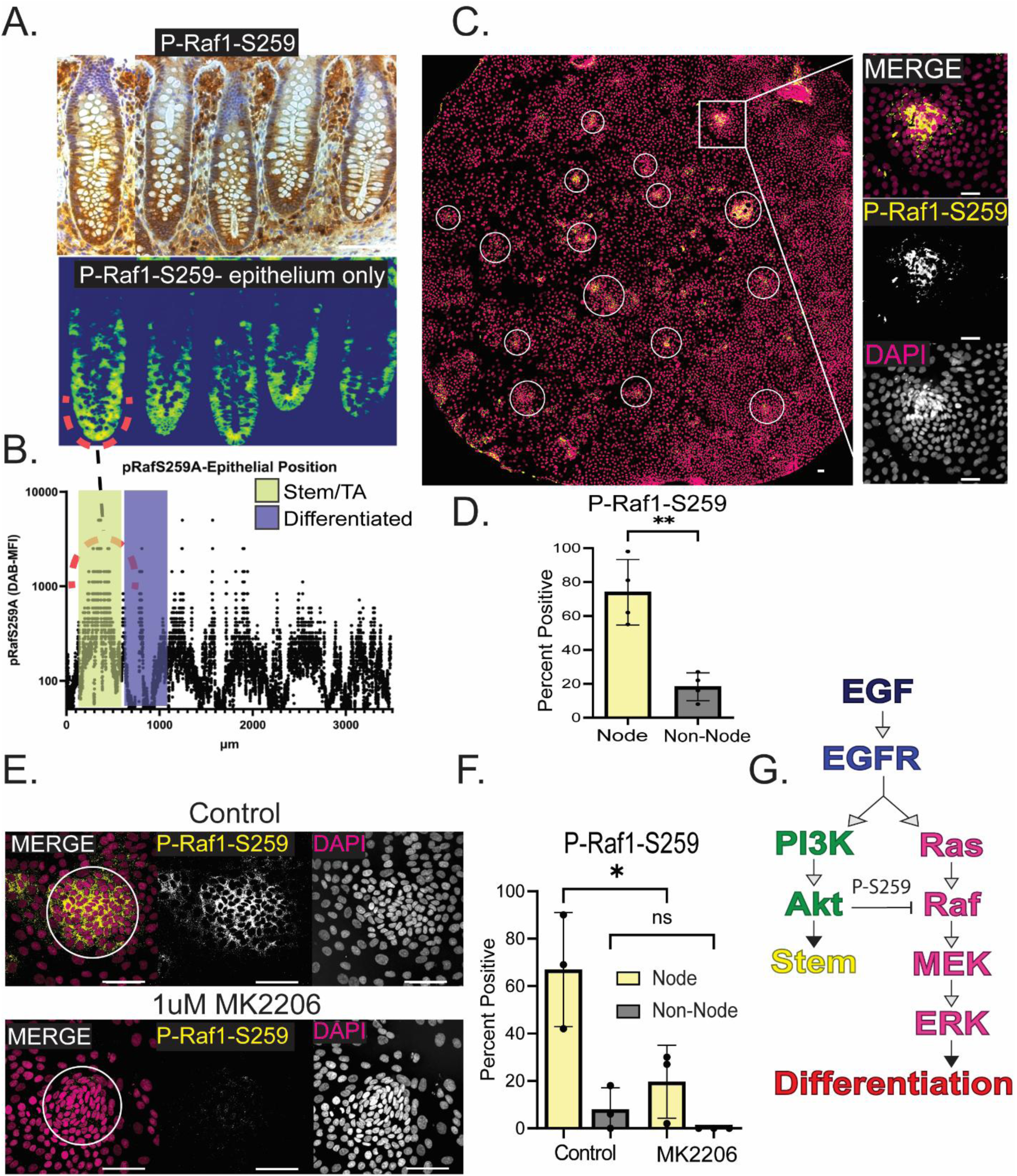
AKT-Dependent RAF-1-S259 Phosphorylation Suppresses ERK Signaling in Stem Cell Niches. **(A)** Representative image of immunohistochemistry stain of P-RAF1-S259 in multiple normal colonic crypts with heatmap showing expression levels in epithelial cells only. **(B)** P-RAF1-S258 IHC quantified as mean fluorescent intensity levels localized to each µm of the top image. Yellow shows stem/TA region as highlighted in the heatmap with a red semicircle, blue shows differentiated region with low phosphorylation. **(C)** Whole 384-well stitch of P-RAF1-S259 immunofluorescent stain in colonic organoids with zoom on one node. **(D)** Quantification of P-RAF1-S259 organoid stain showing activation only within stem cell niches (nodes). **(E)** Representative image of stem cell niche treated with 1µM AKT inhibitor (MK2206) for one hour and control group. **(F)** Quantification of P-RAF1-S259 activity being significantly reduced in nodes after MK2206 treatment. **(G)** Model showing AKT-directed phosphorylation of RAF at serine 259 inhibits MAPK signaling in colonic stem cells. Data shown is from analysis of 4 technical replicates with at least 150 total cells quantified per replicate. Data are represented as mean ± SEM. All scale bars are 100µm. Significance calculated with Welch’s t-test (D) and four-way ANOVA (F), * P ≤ 0.05, ** P ≤ 0.01.

### RAF-1-S259 Functions as an ERK Dynamics Checkpoint

To investigate the potential for AKT to control ERK insulation, we asked if tuning of AKT could regulate ERK signaling dynamics in CSCs. One hour treatment with 1µM MK-2206 significantly reduced the number P-AKT positive cells and overall P-AKT levels in both nodes and non-nodes (Fig. 5A-B, S5B). AKT inhibition also acutely induced activation of ERK specifically within stem cell nodes (Fig. 5C-D, S5C). Similar to PMA treated cells, insulation was preserved (Fig. 5E), ERK dynamics revealed a 2-3 hour significant induction of ERK activity followed by return to baseline levels (Fig. 5F-G), and a significantly increased KDS (Fig. 5H). Loss of AKT activity converted normally insulated stem cells into ERK-responsive cells, resulting in widespread loss of stem cells 72hrs after disruption of the ERK checkpoint without inducing cell death (Fig. 5I-J, S5D-E). Taken together, these data suggest AKT suppresses ERK dynamics within stem cell niches through phosphorylation of RAF-1-S259. When de-repressed, RAF-1 no longer acts as an insulation checkpoint and allows for ERK activation within stem cell niches, leading to global differentiation of the tissue and complete loss of tissue-resident stem cells required for wound-healing (Fig. 5K).

**Figure 5:**
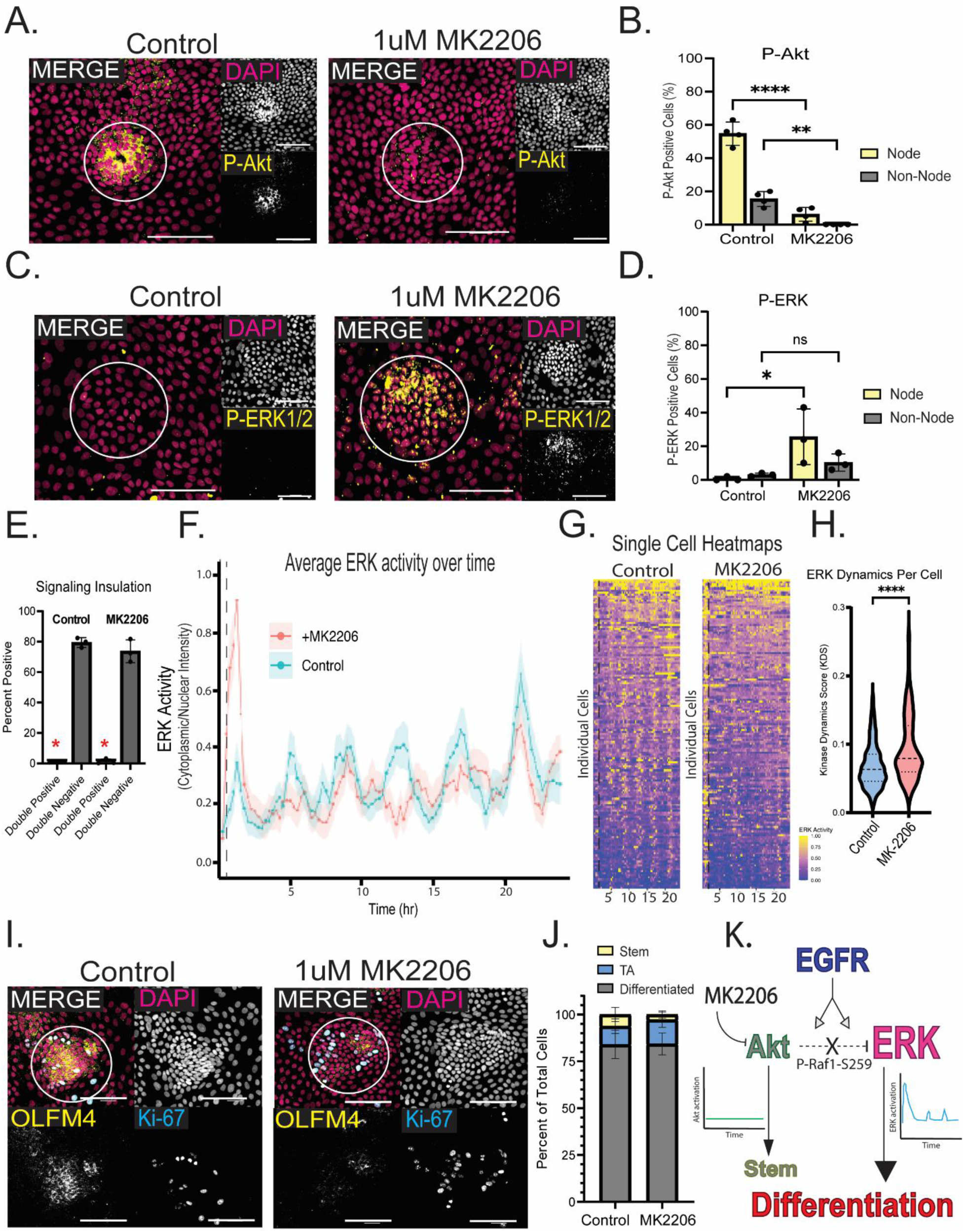
RAF-1-S259 Functions as an ERK Dynamics Checkpoint. **(A)** Control and 1µM MK2206 treated groups representative images of P-AKT staining. **(B)** Quantification of P-AKT in nodes and non-nodes showing significant reduction of AKT signaling throughout entire monolayer. **(C)** Representative image of P-ERK1/2 staining in control and MK2206 treated monolayers. **(D)** Quantification showing significant increase in ERK1/2 signaling specifically within stem cell niches treated with MK2206. **(E)** Quantification of ERK/AKT double positive cells and double negative cells. Red asterisks indicate less than 5% of cells activate both pathways, signaling insulation which is maintained with acute AKT inhibition. **(F)** Quantification of ERK-KTR biosensor in organoids treated with MK2206 and control. Organoids were treated with 1µM MK2206 30 minutes after starting the movie. Graph shows average ERK activity by cytoplasmic/nuclear ration of KTR intensity with ribbon showing 95% CI. **(G)** Heatmaps showing ERK activity of all cells tracked, with each cell being a horizontal line on the graph. Yellow shows high ERK activity and blue being low activity. **(H)** Kinase dynamics score (KDS) violin plot calculated as the standard deviation of ERK activity within each cell over the first 5 hours of the movie. **(I)** Representative image of stem (OLFM4, yellow) and TA (Ki67, blue) cell markers in control and MK2206 treated groups. **(J)** Quantification of the percentage of total cells with stem or TA makers, showing a loss of stem cells and induction of TA cell fate in MK2206 treated group. **(K)** Model showing MK2206 causes a pulse of ERK activity, leading to loss of AKT, loss of stem cells, and global differentiation of the monolayer. Data shown is from analysis of 3-4 technical replicates with at least 150 total cells quantified per replicate. Data are represented as mean ± SEM. All scale bars are 100µm, significance calculated with Welch’s t-test (H) or four-way ANOVA (B,D), * P ≤ 0.05,** P ≤ 0.01, **** P ≤ 0.0001. See also Figure S5, S6, and S7.

To further test the requirement of AKT in suppressing ERK activity to regulate stem cell renewal, we used genetic tuning of AKT signaling. PDCOs overexpressing AKT with a myristylation tag were used to determine the effects of hyperactive-AKT on ERK signaling and stem cell niches. AKT hyperactivity induced significant increases in both AKT and ERK signaling, loss of mutual exclusivity, and a total loss of signaling insulation, with 60% of cell no longer being insulated (Fig. S6 A-B,E). This hyperactivity of both AKT and ERK signaling caused extreme dysplastic-like loss of patterning and node formation, even though RAF1-S259 phosphorylation was increased (Fig. S6 C,F,G). These cells express a constitutively high, unphysiologically relevant levels of AKT. To test how genetically stimulating AKT activation in a temporally controlled model would affect stem cell regulation, we expressed a tamoxifen-inducible AKT-S473D with a myristylation tag system into PDCOs. P-AKT levels were increased with the construct alone but further amplified with the induction of the myristylation (Fig. S7 A-B). Additionally, temporally controlling the induction of AKT allowed for signaling insulation to be maintained, showing a significant decrease in P-ERK while increasing phosphorylation of RAF1-S259 (Fig. S7 C-F). Maintenance of signaling insulation while driving increased AKT activity resulted in an increase in stem and TA population, as expected (Fig. S7 G-H). Importantly, node structure and shape was maintained, indicating a properly patterned monolayer compared to the dysplastic monolayer created by constitutively activating AKT. These data highlight the ability to maintain signaling insulation as a critical requirement for proper stem cell renewal (Fig. S6 H).

These data identify AKT-mediated RAF-S259 phosphorylation as a molecular ERK dynamics checkpoint that buffers stem cells against inappropriate pulsing. Even a single pulse of ERK driven by AKT inhibition or PMA treatment was sufficient to trigger irreversible stem cell differentiation. Additionally, it points to the importance of signaling insulation and proper temporal activation of AKT signaling in maintaining proper tissue patterning.

### Genetic Ablation of ERK Checkpoint Drives a Reversible Neoplastic Mitogenic Signaling State

To test if phosphorylation of RAF-1 by AKT is required to maintain tissue homeostasis through an ERK dynamics checkpoint, we expressed RAF-1-S259A phosphomutant in PDCOs, preventing AKT-mediated inhibitory phosphorylation. In 3D culture, under normal conditions, three-dimensional colonic organoids form a polarized epithelial sphere with a central lumen and multiple budding crypt-like structures (Fig. 6A, left). RAF-1-S259A organoids displayed a dysplastic phenotype with excessive budding and loss of a central lumen, indicative of enhanced proliferation and disorganized growth (Fig. 6A, right). When plated into the 2D monolayer, loss of RAF-1 negative regulation caused a loss of signaling insulation within the organoids, shown by a coactivation of both AKT and ERK, reminiscent of transformed PDCOs (Fig 6B-C, S8A-B). Rather than demonstrating mutual exclusivity, the majority of cells (∼50%) showed loss of signaling insulation via activation of both AKT and ERK (Fig. 6D, S8C). Live ERK dynamics revealed that RAF-1-S259A cells exhibited sustained ERK activation with reduced pulsatility, resulting in a low KDS score, demonstrating an almost complete lack of signaling dynamics in the single cell heatmaps and dynamics analyses (Fig. 6E-G, S8D). RAF-1-S259A displays a dysplastic phenotype which hyperactivates both AKT and ERK while almost completely eliminating the dynamics nature of ERK signaling within the cells (Fig. 6H). In contrast to the ERK checkpoint-deficient RAF-1-S259A, over expression of WT RAF-1 had the opposite effect on cells, driving them into a TA state by partially reducing insulation and KDS, which ultimately lead to loss of stem cells via hyper-differentiation (Fig. S9-10). Taken together these data show that the AKT-dependent ERK checkpoint functions to prevent expansion of neoplastic clones and instead drive ERK active stem cells to terminally differentiate and exit the cell cycle (Fig. 6H).

**Figure 6:**
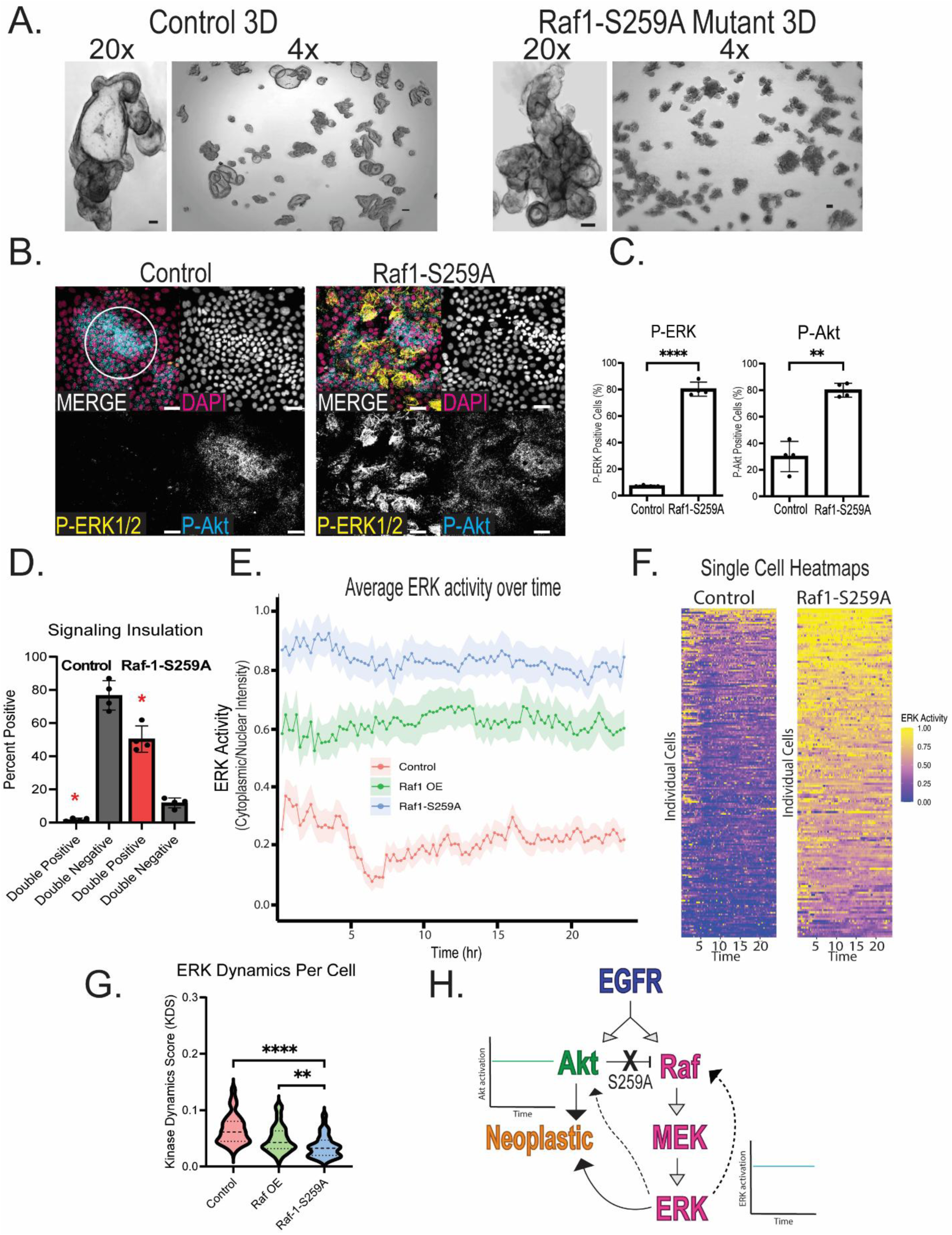
Genetic Ablation of ERK Checkpoint Drives a Reversible Neoplastic Mitogenic Signaling State. **(A)** Representative images of 3D organoids at 4x and 20x zoom. Left shows control organoids with proper lumen and crypt structures, left 3D organoids with RAF1-S259A phosphomutant mutation. **(B)** Representative image of control cells and RAF1-S259 mutant cells stained for P-ERK1/2 (yellow) and P-AKT-473 (blue), showing global upregulation of both pathways. **(C)** Quantification of percent positive of total cells for P-ERK and P-AKT showing a more than 4-fold increase in cells activating each pathway. **(D)** AKT/ERK double negative and double positive quantification, red asterisks highlighting a 10-fold increase uninsulated cells. **(E)** Quantification of ERK-KTR biosensor in control and RAF mutant organoids showing constitutive ERK activation with loss of on/off dynamics. Graph shows average ERK activity by cytoplasmic/nuclear ration of KTR intensity with ribbon showing 95% CI. **(F)** Heatmaps showing ERK activity of all cells tracked, with each cell being a horizontal line on the graph. Yellow shows high ERK activity and blue being low activity. **(G)** Kinase dynamics score (KDS) violin plot calculated as the standard deviation of ERK activity within each cell over the first 5 hours of the movie showing significantly lower ERK dynamics in RAF1-S259A mutant cells than control or RAF overexpression cells. **(E)** Model depicting RAF-S259A phosphomutant causing a global increase in ERK activity that loses negative feedback checkpoint and kinase dynamics inducing high AKT activity, leading to a hyperactive, dysplastic phenotype. Data shown is from analysis of 4 technical replicates with at least 150 total cells quantified per replicate. Data are represented as mean ± SEM. All scale bars are 100µm, significance calculated with Welch’s t-test (C) or by three-way ANOVA (G), ** P ≤ 0.01, **** P ≤ 0.0001. See also Figure S9 and S10.

### Reintroduction of Signaling Dynamics Can Reverse Neoplastic Cell Fate, Regardless of Baseline Signaling Load

Measurement of cell fate in RAF-1-S259A PDCOs showed that loss of signaling insulation and low KDS led to the expansion of a neoplastic, stem-like population that lacked proper tissue patterning or stem cell niche structures (Fig 7A-B). RAF-1-S259A cells lack almost all dynamics within their ERK activity over 24 hours (Fig 7C-E). To test if dynamics is dominant to total kinase activity for driving cell fate decisions, we sought to reintroduce dynamics back into the RAF-1-S259 mutant cells that lacked ERK dynamics. Using both ERK activation (PMA) (Fig. 7C-D) and AKT suppression (MK-2206) (Fig. S11 A-B), we were able to rescue the neoplastic phenotype induced by the RAF-1-S259A mutation by reintroducing a pulse of ERK into the static, non-dynamic cells (Fig. 7 C-E, S11C). Single cell traces and dynamics readouts show a striking induction of dynamics into the RAF-1 mutant lines, even with the elevated baseline ERK signaling in these cells. Importantly, this 4–5-hour pulse of ERK activity (Fig. 7C-purple) was enough to induce a 2-fold decrease in neoplastic cells days later regardless of their baseline ERK signaling load being 3-fold higher compared to controls. Collectively, these findings indicate that the AKT-RAF1-ERK checkpoint is required for homeostasis and prevention of neoplastic transformation by maintaining ERK dynamics in activated stem cells to drive their differentiation. Loss of this dynamics checkpoint can induce hyperactive and dysplastic tissue architecture, reminiscent of early tumorigenesis. This highlights ERK dynamics as epistatic to signaling load in preservation of human CSCs (Fig. 8).

**Figure 7:**
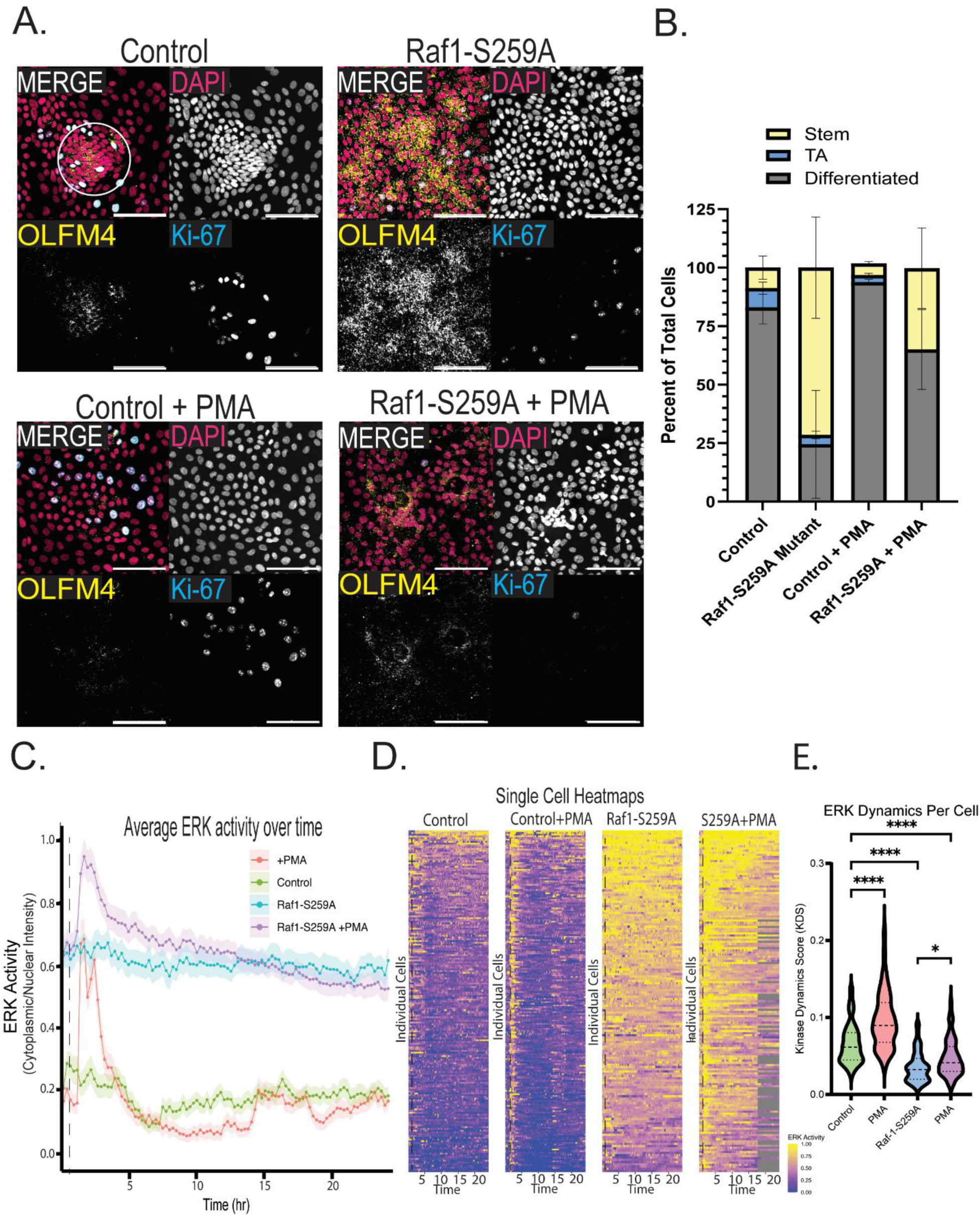
Reintroduction of Signaling Dynamics Can Reverse Neoplastic Cell Fate, Regardless of Baseline Signaling Load. **(A)** Representative images of Control, RAF1-S259A phosphomutant, and both conditions treated with 100nM of PMA for 1 hour then fixed and stained after 72 hours. All wells were stained with OLFM4 (stem cell marker, yellow) and Ki-67 (TA cell marker, blue). **(B)** Quantification of all four conditions showing PMA can rescue the neoplastic cell fate of RAF1-S259A mutant cells. **(C)** Quantification of ERK-KTR biosensor in control and RAF mutant organoids with and without PMA showing PMA can induce a pulse of ERK activity regardless of baseline activity levels. Graph shows average ERK activity by cytoplasmic/nuclear ration of KTR intensity with ribbon showing 95% CI. (**D)** Heatmaps showing ERK activity of all cells tracked, with each cell being a horizontal line on the graph. Yellow shows high ERK activity and blue being low activity. Grey indicates a cell out of focus or field of view. **(E)** Kinase dynamics score (KDS) violin plot calculated as the standard deviation of ERK activity within each cell over the first 5 hours of the movie showing a pulse of PMA is able to induce dynamics back into the RAF1-S259A mutant cells. **(F)** Model showing ERK dynamics regulating cell fate through AKT-dependent RAF1-S259 checkpoint. Breakdown of this checkpoint induced a neoplastic cell fate characterized by high kinase signaling load and low kinase signaling dynamics. Inducing increased kinase signaling dynamics causes global differentiation regardless of kinase signaling load, showing that kinase dynamics are epistatic to signaling load. Data shown is from analysis of 4 technical replicates with at least 150 total cells quantified per replicate. All scale bars are 100uM. Data are represented as mean ± SEM. All scale bars are 100uM, significance calculated with four-way ANOVA, * P ≤ 0.05, ** P ≤ 0.01, *** P ≤ 0.001, ** P ≤ 0.0001. See also Figure S9 and S10. See also Figure S11.

**Figure 8:**
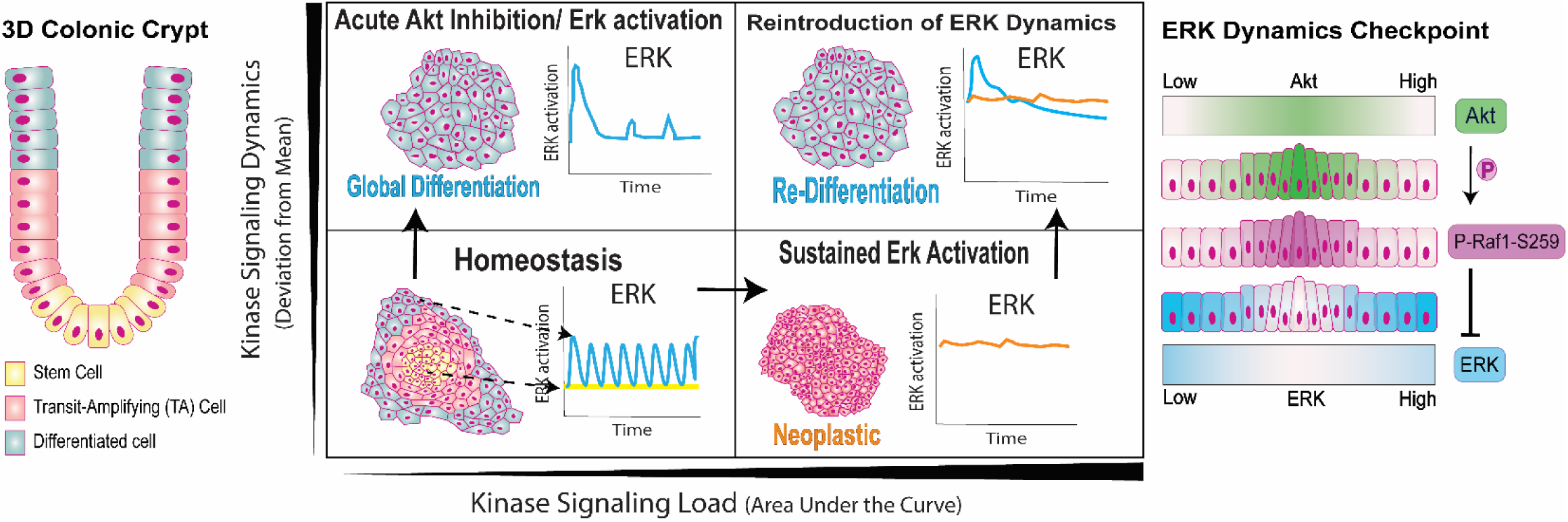
AKT-dependent phosphorylation of RAF-1 at serine 259 establishes a post-receptor checkpoint that maintains ERK temporal dynamics in stem cells. Acute activation of ERK in stem cells triggers rapid global differentiation. Disruption of the ERK checkpoint via mutation of serine 259 leads to sustained AKT and ERK co-activation in stem cells, resulting in the emergence of a neoplastic, architecturally disorganized cell population dominating the cell fate profile. Incredibly, introducing brief ERK pulses through AKT inhibition or ERK activation triggers re-differentiation of neoplastic cells. This highlights ERK dynamics as epistatic to signaling load in preservation of human CSCs.

## DISCUSSION

The intestinal epithelium must preserve a stable stem-cell compartment despite continuous exposure to mitogenic, mechanical, and inflammatory cues. Here, we identify an **AKT-dependent ERK dynamics checkpoint** as a previously unrecognized mechanism essential for preservation of human colonic stem cells. In this model, AKT-dependent phosphorylation of RAF-1 at serine 259 suppresses ERK activation in CSCs despite active EGFR signaling, thereby maintaining a protected stem-cell state within a ligand-rich tissue. Although AKT-mediated inhibition of RAF-1 at S259 has been established biochemically for decades, its role in stem-cell preservation has remained unresolved. This is likely because its function can only be appreciated in systems that preserve long lived CSCs, allow measurement of single-cell kinase dynamics, epithelial architecture, cell-type heterogeneity, and tissue-scale patterning ^25–30^. When this checkpoint is lost, ERK is no longer deployed as a transient lineage-progressive signal, but instead becomes a sustained, non-pulsatile program associated with AKT/ERK co-activation, dysplastic architecture, and hyperactive signaling states.

A central advance of this study is that fate in the human colon is determined not simply by pathway magnitude, but by ERK temporal structure. Prior work established that signaling dynamics can encode distinct cell fate decisions, including transient versus sustained MAPK responses and information transfer through dynamic signaling patterns ^7, 31–35^. Here, brief ERK pulses drive differentiation, whereas sustained ERK without dynamics achieved by oncogenic allele expression, drives the opposite phenotype, a neoplastic state defined by high ERK/AKT/OLFM4, a fate profile almost never observed in our system during homeostasis. Most strikingly, re-introduction of a transient ERK pulse (∼3hrs) is sufficient to re-differentiate cells after 72 hours, even in the setting of oncogenic ERK hyperactivation, demonstrating that ERK signaling dynamics are epistatic to total kinase load. Conceptually, the ERK checkpoint resembles a fold-change-detecting module, buffering absolute mitogenic input while allowing only appropriately structured changes in ERK activity to drive fate transitions ^36^. Our dynamics-based analysis supports this conclusion directly: cell fate segregates more cleanly by dynamic regime than by signal amplitude alone, consistent with the view that temporal organization of ERK activity is a dominant determinant of lineage outcome. This behavior is especially remarkable in light of the classic observation that TPA acts as a tumor promoter rather than an initiator, amplifying growth-promoting programs in susceptible tissues rather than directly establishing transformation^37^. In our system, however, re-introduction of a pulse into hyperactive cells does the opposite of tumor promotion: despite high baseline AKT/ERK signaling, expansion of plasticity markers, and emergence of a dysplastic state, a brief ERK pulse is sufficient to re-drive differentiation.

Our findings also help resolve an apparent paradox in intestinal EGFR signaling. EGFR is broadly required across the epithelium, yet its downstream outputs are not uniformly co-activated. Instead, CSCs sustain AKT while suppressing ERK, whereas differentiated compartments display sporadic AKT/ERK co-activation in wave-like domains associated with epithelial turnover and survival signaling ^8, 19, 30^. We interpret this as a form of signaling insulation, in which shared extracellular input is selectively partitioned into distinct intracellular signaling states. This framework is consistent with broader examples of pathway insulation and biased signaling, including GPCR desensitization, contact-dependent inhibition of EGFR signaling, ligand-specific EGFR dimer stabilization, and non-equivalent Wnt versus R-spondin control of intestinal stem-cell self-renewal ^9, 10, 12, 13, 38, 39^. In this view, the ERK checkpoint is a mechanistic arm of a larger insulation program that allows a continuously stimulated epithelium to preserve stemness while remaining buffered against mitogenic noise.

Several prior observations are clarified by this model. In intestinal organoids, EGFR or MEK/ERK inhibition halts proliferation of Lgr5+ stem cells and induces a reversible quiescent state, supporting the idea that EGFR–ERK signaling contributes to maintenance of proliferative stem-cell output ^40^. Conversely, epithelial ERK1/2 deletion in the developing intestine causes expansion of stem-like compartments, disruption of the normal Ki67+ progenitor architecture, and defective differentiation, consistent with the idea that ERK is required not merely for proliferation, but for orderly progression out of the stem-cell state ^41^. Together with our data, these studies argue that the critical variable is not whether ERK is simply on or off, but how, where, and for how long ERK is activated. We propose that pulsatile ERK promotes CSC exit into a transient TA state and subsequent differentiation, whereas sustained activation destabilizes fate control and promotes formation of neoplastic cell lineages. Mechanistically, RAF-S259A likely abolishes ERK dynamics not because S259 itself is the sole feedback mechanism for recovery of basal ERK levels, but because it removes a major inhibitory gate on RAF membrane recruitment and Ras coupling, preventing efficient re-establishment of the autoinhibited state after activation ^42–44^. In that light, the limited effect of gefitinib in RAF-S259A cells is expected: once signaling has shifted into a receptor-bypassing RAF-driven state, EGFR inhibition can no longer efficiently extinguish downstream pathway activity.

An important implication is that signaling insulation in CSCs is likely established by multiple upstream layers rather than RAF-S259 alone. EGFR ligands can impose distinct signaling kinetics through differential receptor dimer stabilization, receptor output can be attenuated by contact-dependent inhibitory mechanisms, and tissue-scale properties such as density, cell shape, compression, and matrix mechanics can pre-pattern heterogeneous cell states and bias fate transitions ^9, 45–49^. These studies support a model in which CSCs do not passively interpret ligand abundance but actively maintain an insulated state through coordinated control of receptor output, mechanics, and extracellular context. In that framework, the extracellular matrix may function as an innate insulation layer that helps preserve stem-cell identity by stabilizing the physical and signaling environment in which AKT remains high and ERK remains suppressed.

This model also helps reconcile homeostasis with regeneration. The intestine must preserve stem-cell fidelity over long timescales while remaining capable of rapid repair after injury, and increasing evidence suggests that this regenerative capacity is distributed across a broader plastic epithelial system rather than a single dominant progenitor ^50–52^. We therefore favor a model in which self-organization first establishes a patterned epithelial architecture, after which signaling insulation preserves that architecture by partitioning downstream EGFR outputs across cell states. Within this hierarchy, the ERK checkpoint represents one essential insulation layer that protects CSCs from inappropriate differentiation while preserving the capacity to deploy ERK transiently for migration, survival, and repair. More broadly, our findings highlight the need to measure signaling at the single-cell dynamic level in organoid and tissue-scale systems, because only these approaches can capture how cellular communication is partitioned across heterogeneous epithelia and how native epithelial mechanisms preserve homeostasis in the gut and its associated pathologies. We propose that preservation of human colonic stem cells depends on a hierarchically organized insulation program in which self-organization, extracellular context, and post-receptor signaling checkpoints cooperate to suppress inappropriate ERK activation, thereby allowing a continuously stimulated epithelium to remain both stable and regenerative.

## LIMITATIONS OF THE STUDY

Pharmacological perturbations of AKT and ERK are useful to induce transient signaling changes, but has the potential for off target effects. The *in-vivo* stem cell niche is exceptionally difficult to fully replicate *ex-vivo* as many coordinated inputs are at work. Here, we study an innate regulatory mechanism of adult colonic stem cells, however proper tissue function relies on coordinated signaling across cell types and scales. Intravital imaging and alternative methods of studying single stem cell signaling dynamics in adult colonic tissues has not been developed, and would enhance the model presented here.

## RESOURCE AVAILABILITY

### Lead contact

Further information and requests for resources and reagents should be directed to and will be fulfilled by the lead contact, Kelvin Pond

### Materials availability

All the materials generated in this study are available upon request from the lead contact. This study did not generate unique reagents.

### Data and code availability

All data reported in this study are available from the lead contact upon request. This paper does not report the original codes. Any additional information required to reanalyze the data reported in this paper is available from the lead upon request.

## ACKNOWLEDGMENTS

We thank members of the Pond and Thorne labs for helpful comments and discussion. This work was supported by Colorectal Cancer Alliance Young Investigator Award 10041525 (K.W.P), and National Institutes of Health grants R01DK137411, R35GM147128 (C.A.T.). Tissue acquisition was supported by the TACMASR core at University of Arizona, grant P30 CA023074. Imaging was performed in part through the Nikon Center of Excellence at the University of Arizona Cancer Center. Flow cytometry sorting was supported by the University of Arizona Flow Cytometry and Human Immune Monitoring Shared Resource, grant CCSG - CA 023074.

## AUTHOR CONTRIBUTIONS

Conceptualization and experimental design, L.R., A.B., K.P.; methodology and data acquisition, L.R., A.B., J.C.; data analysis and figure preparation, L.R., S.M., S.M., K.W.P.; manuscript writing, review, and editing, L.R., A.B., C.A.T., K.W.P.; project supervision, C.A.T., K.W.P.

## MATERIALS AND METHODS

### Human specimens and organoid derivation

Normal and tumor colonic tissues were obtained from consented patients undergoing endoscopic ultrasound-guided fine-needle aspiration or core needle biopsy under protocols approved by the University of Arizona IRB (TARGHETS, IRB 1909985869). Biospecimens were deidentified by the Tissue Acquisition and Cellular/Molecular Analysis Shared Resource (TACMASR) prior to distribution. Investigators had no access to identifiable information. The study was therefore exempt from human subject classification.

Biopsy specimens were transported in Advanced DMEM/F12 supplemented with GlutaMax, HEPES, Amphotericin B, Gentamycin, Kanamycin, N2, B27 minus vitamin A, N-acetylcysteine, nicotinamide, CHIR99021, and Thiazovivin. Tissues were minced, cryopreserved in 70% seeding media supplemented with 20% FBS and 10% DMSO, and stored at the BioDROids core facility. For organoid establishment, thawed tissue fragments were digested with 1 mg/ml collagenase type III for 10 to 25 minutes at room temperature, washed in PBS, embedded in 100% Matrigel, and cultured in seeding media for 7 days. Low-passage aliquots were cryopreserved or used for lentiviral infection.

### Organoid culture conditions

Organoids were maintained in Matrigel (Corning, 356231) with complete LWRN medium consisting of Advanced DMEM/F12 (Gibco, 12634-010) supplemented with 1x GlutaMax (Gibco, 35050-061),10nM HEPES (Corning, 25-060-CI), 1x N2-MAX (R&D Systems, AR009), 1x N21-MAX (R&D Systems, AR008), 1mM N-acetylcysteine (Sigma-Aldrich, A8199-10G), 1% penicillin/streptomycin (Gibco, 15070063), 50% L-WRN conditioned mediµM,100ng/mL recombinant human EGF (PeproTech, AF-100-15-500UG), 10 µM SB202190 (Stemcell Technologies, 72632), and 1x Primocin (InvivoGen, ant-pm-1).

Seeding media consisted of complete LWRN supplemented with 2.5 µM CHIR99021 (Sigma-Aldrich, SML1046-5MG) and 10 µM Y27632 (Tocris, 1254).

### Lentiviral transduction

HEK293T cells were maintained in DMEM (Corning, 10-013-CV) supplemented with 10% FBS and 1% penicillin/streptomycin (Gibco, 15070063) and plated to obtain 40% confluence for transfection. 3.25µg psPAX2, 1.75µg pMD2.G, and 5µg plasmid of interest were combined and transfected into HEK293T cells using GeneJuice according to the manufacturer’s recommendations. Media was replaced after 16hrs of transfection and viral supernatant was collected every 24hrs for 2 days, storing at 4°C. Viral supernatant was centrifuged at 1600 x g for 10 minutes to pellet cell debris and filtered through a 0.45µm syringe filter. Cleared virus was concentrated overnight at 4°C using LentiX concentrator (Takara Bio, 631231) according to the manufacturer’s instructions. C Viral supernatant was concentrated 100-fold using LentiX concentrator (Takara Bio, 631231) concentrated viral pellets were resuspended in organoid seeding media and stored at −80°C until use.

For infection, organoids were harvested and dissociated into single-cells as described in “2D preparation” section. Dissociated organoid suspensions were mixed 1:1 with an equal volume of concentrated viral stock supplemented with 5µg/mL polybrene (VectorBuilder, PL001) in a 48-well plate (Fisher, FB012930). organoid cells were resuspended in viral media containing polybrene, Organoid viral mix was centrifuged at 600 x g for 1 hour at 32°C and then incubated at 37°C for 3 hours. Cells were harvested by manual pipetting, centrifuged at 500 x g for 5 minutes, and re-embedded in 90% Matrigel. Matrigel was overlayed with seeding media for 24hrs after viral infection. Seeding media was replaced with and maintained in complete LWRN until fully grown. Infection efficiency exceeded 50% prior to downstream experiments. Dual-reporter organoids were FACS sorted using a BD FACS Aria III to normalize reporter expression levels. For RAF OE, Myr-AKT, and RAF-S259A, 2µg/mL puromycin was added complete LWRN for 3 days to select for cells containing the construct.

For all ERK-KTR dynamics figures, RAF OE, and constitute Myr-AKT, infection was done as cells were being prepared for 2D monolayers, rather than grown as 3D organoids then plated into 2D monolayers due to difficulties with passaging in 3D. Cells were prepared for 2D monolayers as below, and viral media was added to seeding media prior to plating into 384-well plate. Lentivirus was added at a 1:10 concentration for Myr-AKT construct and 3:4 concentration for RAF-S259A and RAF overexpression constructs. Plate was centrifuged at 600 g for 1 hour at 32°C, stored in 37°C for 1 hour. Wells were then washed with complete media three times before being stored in seeding media overnight. After 24 hours, 2ug/mL puromycin was added complete LWRN for 2 days to select for cells containing the construct.

### Preparation of 2D organoid monolayers

Organoids were harvested from Matrigel using ice-cold Advanced DMEM/F12 (Gibco, 12634-010) and manual pipetting. Organoid suspension was kept cold and centrifuged at 500 x g for 5 minutes. The supernatant was removed and three additional washes with Advanced DMEM/F12 were performed to separate organoids from Matrigel. After washes, organoid pellets were suspended in 0.25% Trypsin (Corning, 25-053-CI) supplemented with 10µM Y27632 and warmed to 37°C for 5 minutes to dissociate. Trypsin was quenched with an equal volume of FBS and dissociated organoids were pelleted at 500 x g for 5 minutes. Supernatant was removed and pellets were resuspended in seeding media. Dissociated organoids were plated at 7000 cells per well into 384-well plates (Greiner, 781866) that were pre-coated with a 1:40 dilution of Matrigel in Advanced DMEM/F12 for 1 hour at 37°C. Seeding media was removed after 24hrs and replaced with complete LWRN. Media was changed daily as monolayers became confluent and spatial patterning emerged, typically after 5-7 days.

### Live-cell ERK dynamics imaging

ERK activity was monitored using ERK-KTRmRuby2 with H2B-iRFP670 nuclear segmentation. Time-lapse imaging was performed on a Nikon Eclipse Ti2 automated microscope at 15-minute intervals for 24 hours.

Nuclei were segmented using custom MATLAB scripts based on H2B. Cytoplasmic ERK-KTR intensity was calculated using a two-pixel annulus surrounding each nucleus. ERK activity was defined as cytoplasmic intensity exceeding nuclear intensity.

### Quantification of ERK dynamics states

Tracking data was processed and figures were generated in R 4.5.3 using various packages including tidyverse 2.0.0, readxl 1.4.5, ggplot2 4.0.2, pheatmap 1.0.13, confintr 1.0.2, and RColorBrewer 1.1-3. Tracked nuclear / cytoplasmic activity ratios were inverted to convert them to cytoplasmic / nuclear activity ratios. Raw data was then normalized within each set of conditions by scaling individual cell activity values to a range determined by the global minimum activity and global maximum activity, such that the final activity ratios resided within a 0 to 1 range. Population traces were generated by computing the mean of all cells in each dataset at each time point along with a 95% confidence interval, and plotted using ggplot2’s geom_line() function for the mean, and geom_ribbon for the confidence intervals. Heatmaps were generated using pheatmap on the normalized activity data for each cell within each dataset.

Kinase dynamics analysis was performed on raw nuclear / cytoplasmic activity ratios. Each cell standard deviation was calculated for the first 5 hours of the movie and averaged, then plotted as a single point along a violin plot in Graphpad Prism.

### EGF pulse

HCT-8 colonic adenocarcinoma cell line and HCEC colonic immortalized cell lines were maintained in DMEM (Corning, 10-013-CV) supplemented with 10% FBS and 1% penicillin/streptomycin (Gibco, 15070063). Cells were plated in a 96 well plate and grown until confluent along with PDCOs. All cells were starved from EGF and serµM for 24 hours. 100ng/mL EGF was added to each cell line 10 minutes after movie start. Images of ERK-KTRmRuby2 with H2B-iRFP670 were taken every 5 minutes for 80 minutes. Average ERK activity as determined as above.

### Pharmacologic perturbations

Confluent monolayers were treated with:

- MK-2206 (AKT inhibitor) (Selleck Chemicals, S1078) (1uM)
- PMA (MAPK activator) (Cell Signaling Technology, 4174S) (100nM)
- Gefitinib (EGFR inhibitor) (MedChemExpress, HY-50895) (500nM)
- 4-Hydroxytamoxifen (MedChemExpress, HY-16950) (1uM)

Short-term signaling experiments were performed with 1-hour treatment prior to fixation. Fate transitions were assessed after 72 hours of continuous exposure for MK-1106 or 1 hour pulse-wash for PMA.

### Immunofluorescence

Media was removed and monolayers were immediately fixed in paraformaldehyde (Electron Microscopy Sciences, 15710) diluted to 4% in DPBS (Corning, 21-031-CV) for 7 minutes at room temperature. Monolayers were washed three times with DPBS after fixation and permeabilized with 0.2% Triton X-100 (Sigma, X100-100ML) in DPBS for 10 minutes at room temperature. Monolayers were washed three times with DPBS after permeabilization and blocked with 2.5% BSA (Jackson, 001-000-162) for 1 hour at room temperature. Primary antibodies were diluted in 2.5% BSA and monolayers were incubated with primaries overnight at 4°C. After primary antibody incubation, monolayers were allowed to warm to room temperature and washed three times with 0.1% Tween-20 (Sigma, P9416-100ML) diluted in DPBS. Secondary antibodies and DAPI (Sigma, D9542) were diluted in 2.5% BSA and applied for 2 hours at room temperature. Monolayers were washed three times with 0.1% Tween-20 and stored in DBPS before imaging. Whole-well z-stacks were acquired on a Nikon SoRa spinning disk confocal microscope and projected using extended depth-of-focus reconstruction.

Analysis of immunofluorescent images were done using Nikon General Analysis 3 program. Cells were individually segmented with DAPI brightness and grown to a 20uM diameter. Antibody signal was segmented individually for each antibody and each experiment. Percent positive quantifications were performed by quantifying percent positive for each antibody divided by total cells. Nodes were segmented by DAPI brightness, as node clusters are brighter due to higher cell density.

### Immunohistochemistry

Leica Bond RXm automatic immunohistochemistry/in-situ hybridization platform was used. Staining was performed using the Bond Polymer Refine Detection Kit (Leica Biosystems, DS8900) following the standard “IHC Protocol F” staining protocol for mouse and rabbit primary antibodies. Images were taken using Nikon Eclipse Ti2 microscope.

### EdU incorporation and proliferation analysis

EdU labeling was performed using Click-iT EdU imaging kits according to manufacturer protocols. Ki67 staining was used to identify proliferative compartments. Quantification was performed using automated segmentation pipelines.

### Statistical analysis

All experiments were performed with 3-10 independent biological replicates unless otherwise stated. Data are presented as mean ± SEM. Statistical significance was determined using two-tailed t-tests or ANOVA with appropriate post hoc correction in GraphPad Prism. Exact n values and statistical tests are provided in figure legends. All statistical analyses were performed with GraphPad Prism software.

### Plasmids and Antibodies

pLentiPGK DEST H2B-iRFP670 was a gift from Markus Covert, addgene #90237 (Nguyen et al., 2015). pLentiPGK BLASTDEST ERK-KTRmRuby2 was a gift from Markus Covert, addgene #90231 (Nguyen et al., 2015). pLenti-puro/RAF1-S259A was a gift from Jacques De Grève (Addgene plasmid # 131727, Noeparast et al.,2019[LR9.1]). pLenti-puro/RAF1 was a gift from Jacques De Grève (Addgene plasmid # 131725, Noeparast et al.,2019). Myr-Akt constitutive and Myr-Akt-473D vectors were a generous gift from John Albeck^53^. Antibodies and dyes Rabbit monoclonal OLFM4 (CST, 14369S), Rat monoclonal anti-Ki67 (Invitrogen, 14-5698-82), Mouse monoclonal P-ERK1/2 (Invitrogen, 14-9109-82), Rabbit Monoclonal P-AKT-Ser473 (CST, 4058), Rabbit monoclonal P-RAF1-S259 (Invitrogen, 44-502), Rabbit monoclonal P-EGFR-Y1068 (abcam, ab40815), Mouse monoclonal EGFR (Invitrogen, MA5-13070), Click-iT EdU Cell Proliferation Kit for Imaging, DAPI (Thermofisher, D21490), Goat anti-Rabbit IgG (H+L) Highly Cross-Adsorbed Secondary Antibody, Alexa Fluor Plus 488 (Invitrogen, A32731), Goat anti-Rat IgG (H+L) Highly Cross-Adsorbed Secondary Antibody, Alexa Fluor Plus 647 (Invitrogen, A21247), Goat anti-Mouse IgG (H+L) Highly Cross-Adsorbed Secondary Antibody, Alexa Fluor Plus 647 (Invitrogen, A32728), Goat anti-Mouse IgG (H+L) Highly Cross-Adsorbed Secondary Antibody, Alexa Fluor Plus 546 (Invitrogen, A11030).

