## Supplemental Data for "Preservation of Human Colonic Stem Cells Requires an ERK Dynamics Checkpoint Mediated by AKT"

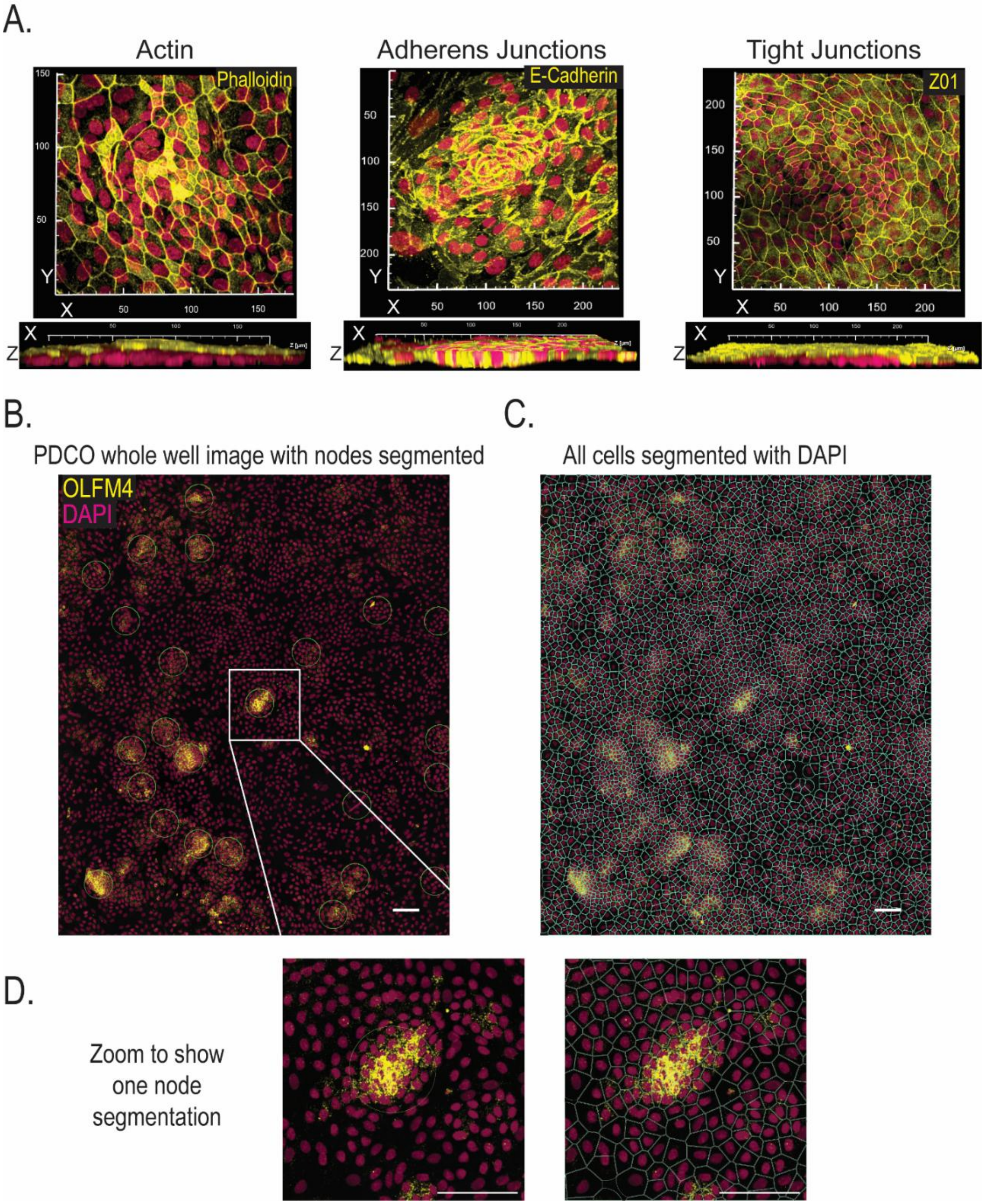


**Figure Supplement 1: PDCO self-organized into polarized and quantifiable monolayers**, related to figure 1

**(A)** Representative image of Actin through staining for phalloidin, adherens junctions shown by staining for E-cadherin, and tight junctions through staining for Z01 in PDCOs. Tight junctions localize on top of nuclei with adherens junctions in between cells, highlighting correct cell orientation within monolayer. **(B)**

Representative image of whole 384-well plate stitch of 2D organoid monolayer stained with DAPI and OLFM4, with nodes segmented in green through DAPI brightness. **(C)** Representative image of all cells in monolayer segmented. Individual cells are segmented using DAPI and have grown 20µm to fill cell area. **(D)** Representative image of zoomed image of one node segmented for node and segmented for all cells. All scale bars are 100µm.

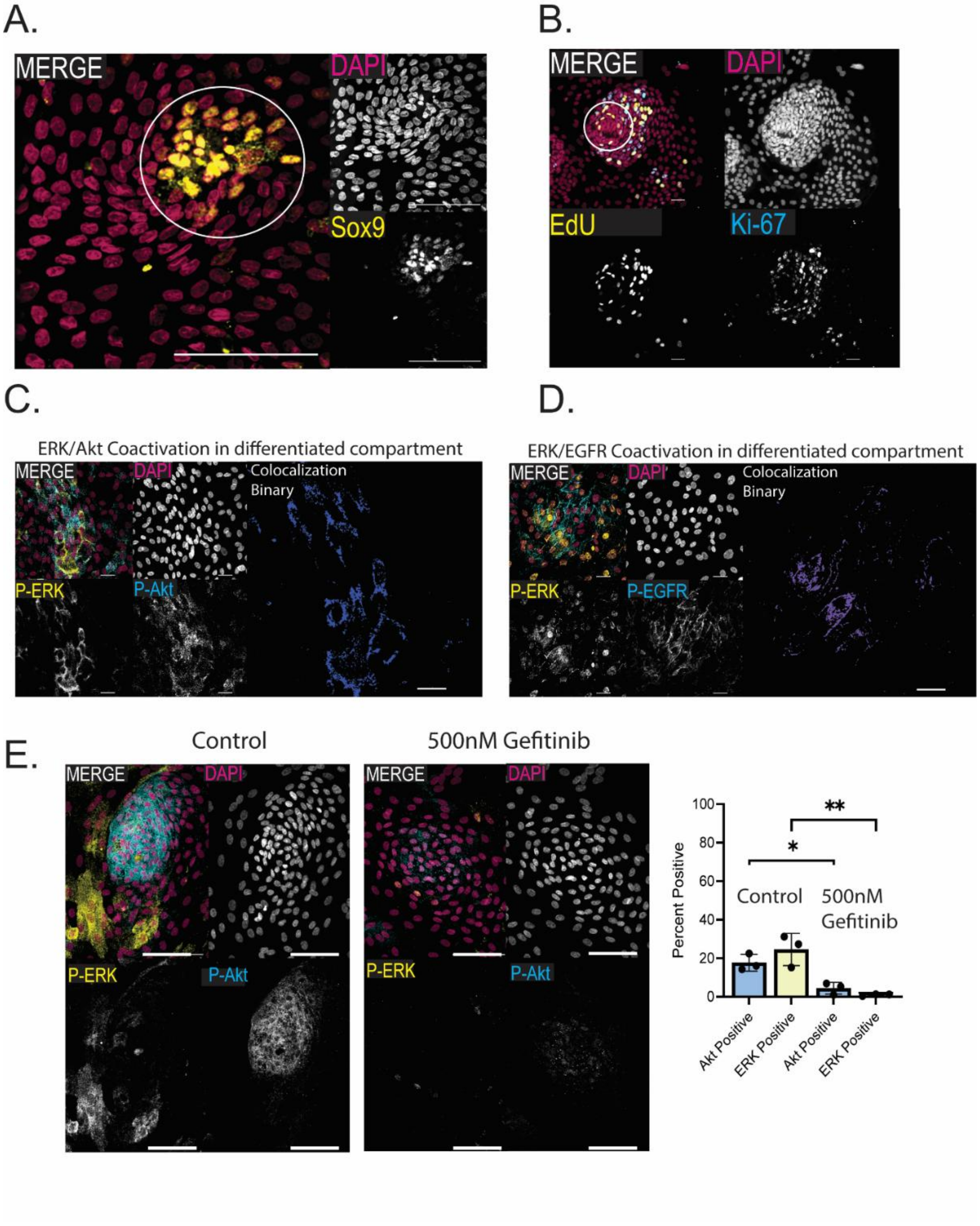


**Figure Supplement 2: Colonic stem cells maintain unique signaling compared to differentiated compartment despite mutual dependence on EGFR**, related to Figure 1

**(A)** Representative image of Sox9 showing activation only in stem cell niches. **(B)** Representative image of Edu (yellow) and Ki-67 (blue). Both show proliferation primarily in and directly around stem cell niches. **(C)** Co-stain of P-ERK1/2 (yellow) and P-Akt-S473 (blue) within differentiated cells undergoing mechanical tension, likely following apoptotic event in middle. **(D)** Co-stain of P-ERK1/2 (yellow) and P-EGFR-Y1068 (blue) showing co-activation in differentiated cells. Both are rare events usually following apoptosis of a nearby cell creating a wave of activation in uninsulated differentiated cells. **(E)** Representative images and quantification of P-ERK and P-Akt inactivation due to ERGF inhibition by 500nM Gifitinib over 1 hour. Quantification is of triplicate technical replicates with at least 150 total cells quantified per replicate. All scale bars are 100µm. Data are represented as mean ± SEM. significance calculated with four-way ANOVA, * P ≤ 0.05, ** P ≤ 0.01.

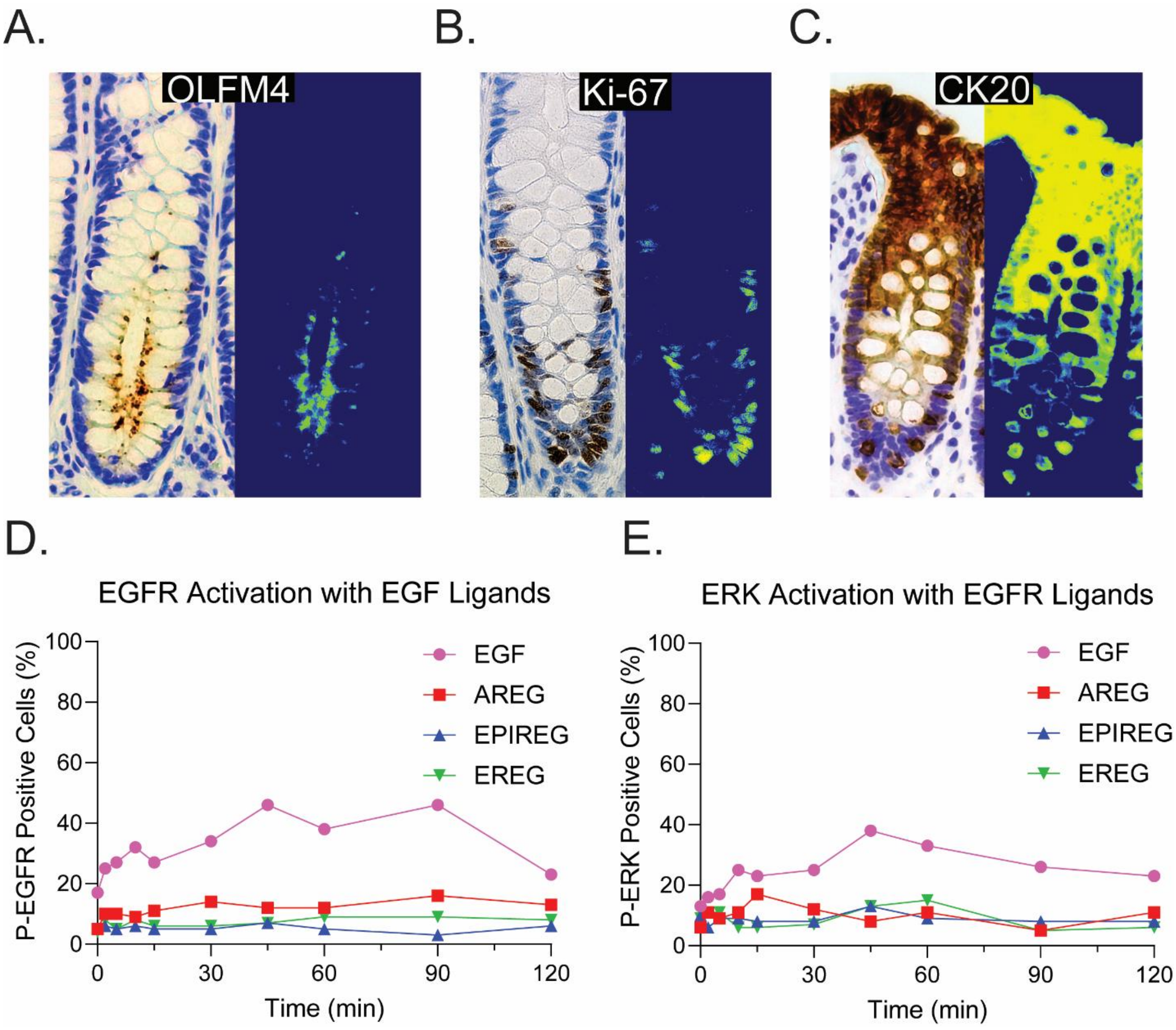


**Figure Supplement 3**: **PDCOs recapitulate *in vivo* patterning but do not respond uniformly to EGFR ligands**, related to Figure 2

**(A)** Representative image of OLFM4 immunohistochemistry staining in normal human colonic tissue showing stem cells at the crypt base. **(B)** Representative image of Ki-67 immunohistochemistry staining in normal human colonic tissue showing the transit-amplifying cell region. **(C)** Representative image of CK20 immunohistochemistry staining in normal human colonic tissue showing differentiated cells at the top of the crypt. **(D)** Quantification of P-EGFR-Y1068 activation with 100ng/mL of various EGFR ligands. EGF shows activation in around 50% of cells at 45 minutes with the rest showing little to no activity. **(E)** Quantification of P-ERK1/2 activation with 100ng/mL of various EGFR ligands. EGF shows activation in around 40% of cells at 45 minutes with the rest showing little to no activity. Three to four biological replicates were performed. All scale bars are 100μm.

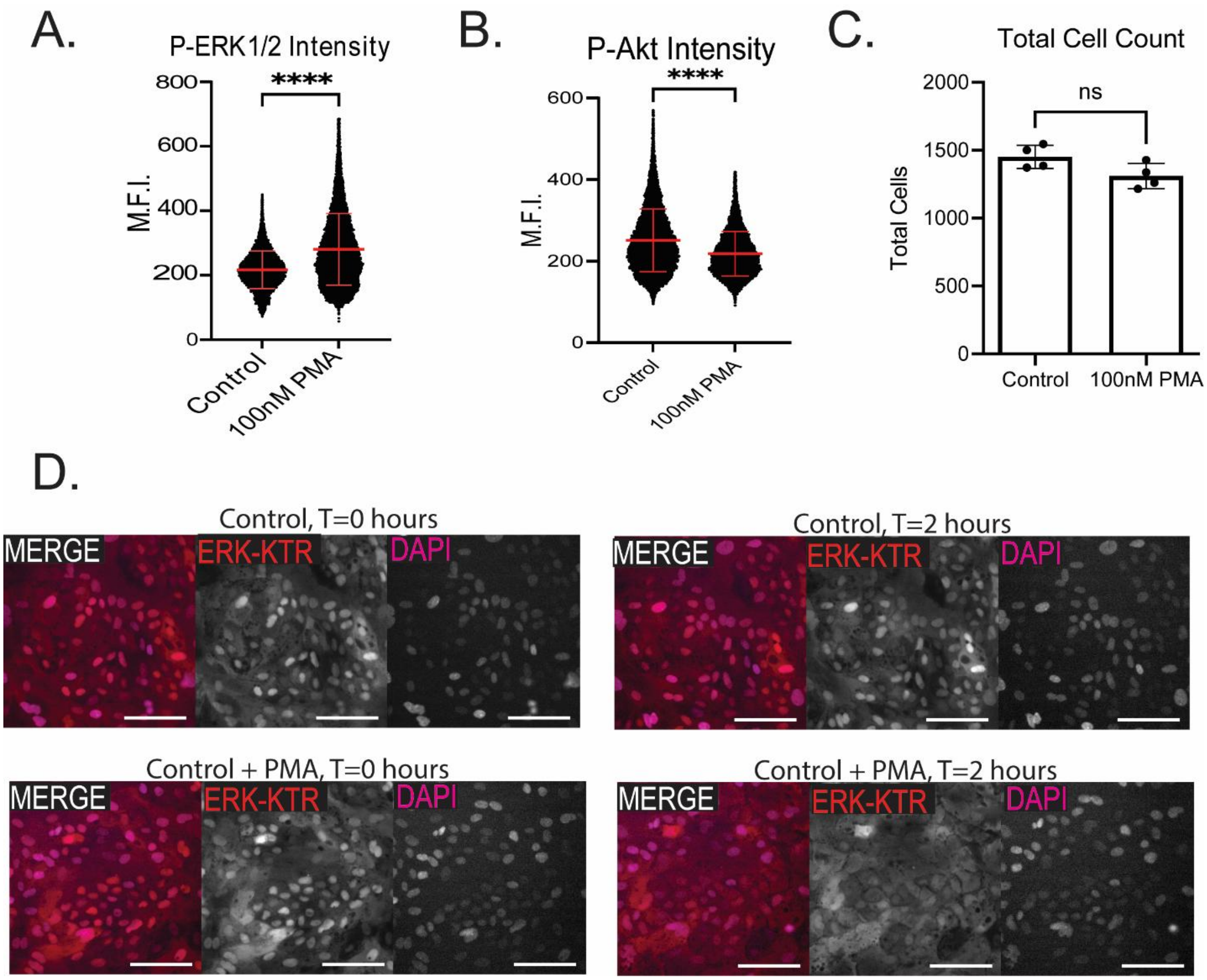


**Figure Supplement 4**: **A single pulse of ERK reduces AKT signaling without increasing cell death**, related to Figure 3

(**A**) Quantification of P-ERK1/2 intensity in control and PMA treated wells, shown as the mean fluorescent intensity (MFI) of each cell in the monolayer. **(B)** MFI quantification of P-Akt within the monolayer 1 hour after PMA treatment or control. **(C)** Total amount of cells in the PMA treated versus control groups showing no change in cell number indicating large amounts of death within the culture. **(D)** Representative images of ERK-KTR organoids in control groups or PMA groups 2 hours after treatment. PMA groups show global EKR activation whereas control shows minimal change in activity. MFI quantifications are of hundreds of cells within one biological replicate, at least three total biological replicates were performed. Three to four biological replicates were performed. Data shown is from analysis of 4 technical replicates with at least 150 total cells quantified per replicate. Data are represented as mean ± SEM. All scale bars are 100μm, significance calculated with ANOVA test, **** P ≤ 0.0001

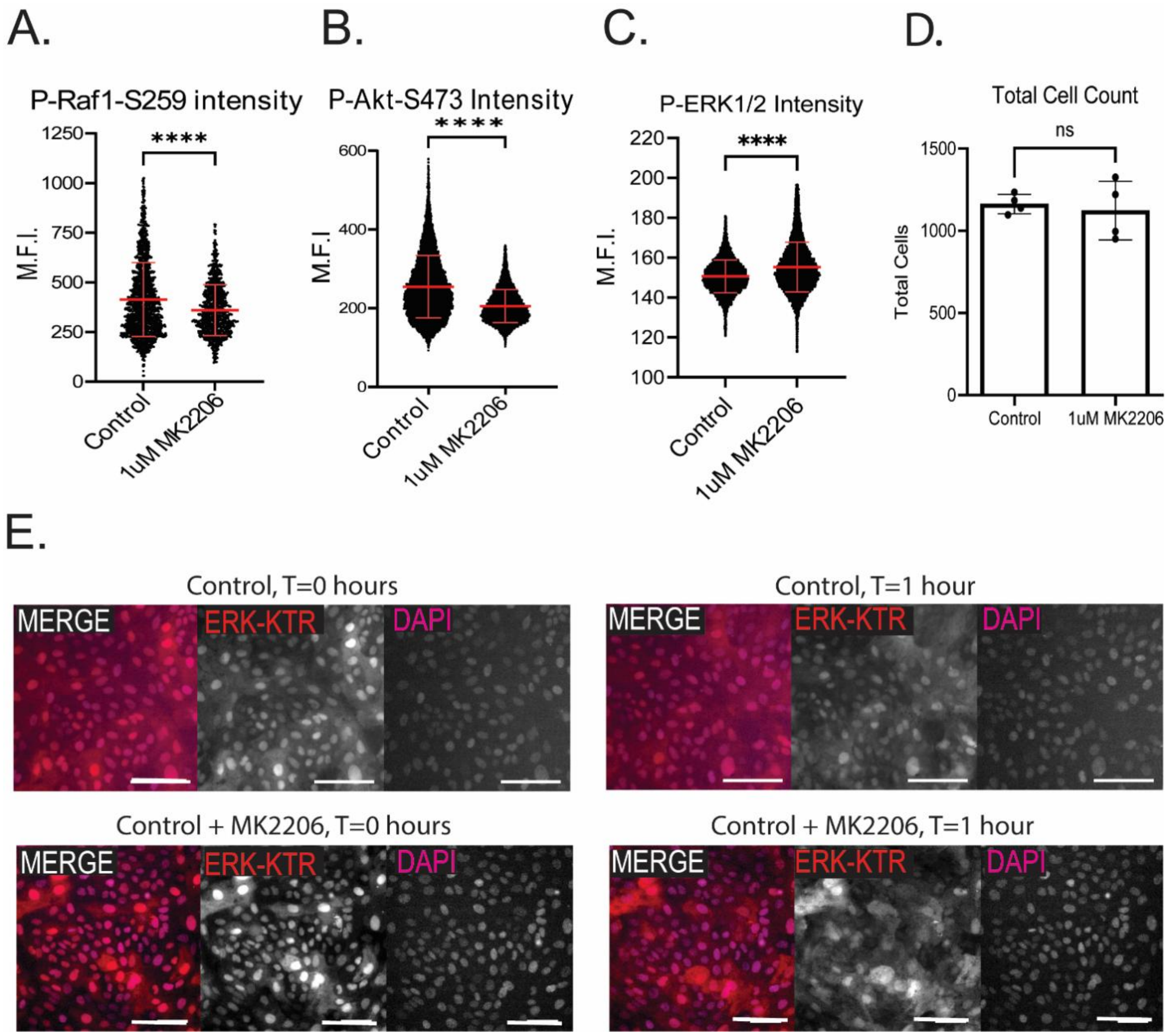


**Figure Supplement 5**: **AKT inhibition induces ERK pulse without cell death**, related to Figure 5

**(A)** Quantification of P-Raf1-S259 intensity in control and 1μM MK2206 treated wells, shown as the mean fluorescent intensity (MFI) of each cell in the monolayer. **(B)** Quantification P-Akt-S473 intensity in control and 1μM MK2206 treated wells, shown as the mean fluorescent intensity (MFI) of each cell in the monolayer. **(C)** Quantification of P-ERK1/2 in control and 1μM MK2206 treated wells, shown as the MFI of each cell in the monolayer. **(D)** Total amount of cells in the MK2206 treated versus control groups showing no change in cell number indicating large amounts of death within the culture. **(E)** Representative images of ERK-KTR organoids in control groups or MK2206 group 30 minutes after treatment. MK2206 groups show global ERK activation whereas control shows minimal change in activity. MFI quantifications are of hundreds of cells within one biological replicate, at least three total biological replicates were performed. Data are represented as mean ± SEM All scale bars are 100μm, significance calculated with ANOVA test, **** $P \leq 0.0001$.

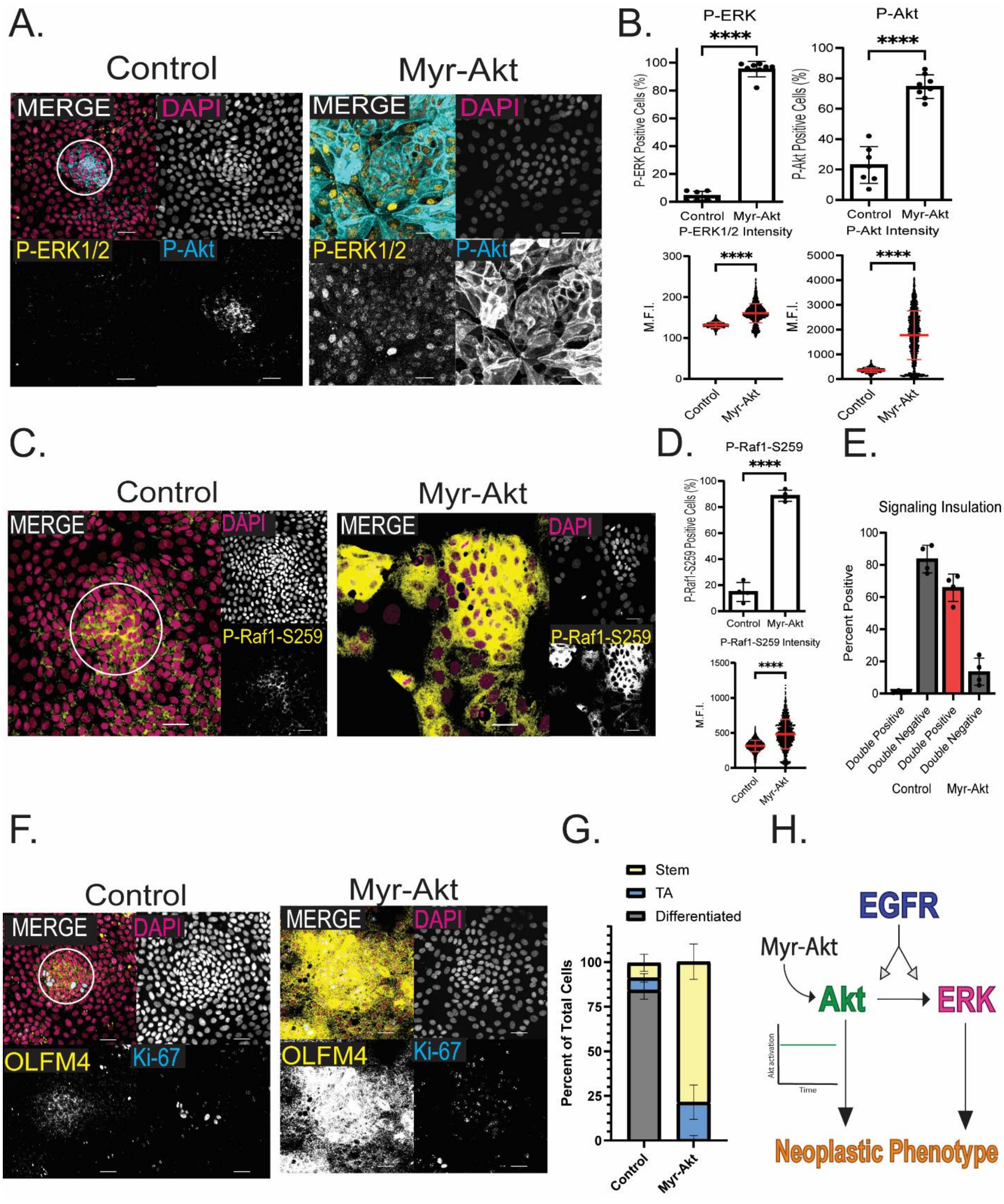
A.
Control
Myr-Akt
MERGE
DAPI
P-ERK1/2
P-Akt
B.
P-ERK
P-Akt
****
P-ERK Positive Cells (%)
P-Akt Positive Cells (%)
Control
Myr-Akt
P-ERK1/2 Intensity
P-Akt Intensity
M.F.I.
C.
D.
P-Raf1-S259
P-Raf1-S259 Positive Cells (%)
P-Raf1-S259 Intensity
E.
Signaling Insulation
Percent Positive
Double Positive
Double Negative
F.
OLFM4
Ki-67
G.
Stem
TA
Differentiated
Percent of Total Cells
H.
EGFR
Myr-Akt
Akt
ERK
Akt activation
Time
Neoplastic Phenotype

**Figure Supplement 6: AKT negative feedback is required for signaling insulation**, related to Figure 5

**(A)** Representative image of control cells and cells infected with lentiviral myristoylated Akt construct causes constitutive hyperactivity of Akt. Staining for P-ERK1/2 (yellow) and P-Akt (blue) shows extreme activity of Akt. P-Akt stain lookup table (LUTs) are not matched due to extreme brightness of Myr-Akt antibody stain. P-ERK stain is matched. Lentivirus was added to single cells during monolayer preparation due to difficulties growing the organoids in 3D. **(B)** Quantification of percent positive of total cells for P-ERK and P-Akt showing significant hyperactivation of both pathways and mean fluorescent intensity (MFI) of each cell within the monolayer for P-ERK and P-Akt activation. **(C)** Control and Myr-Akt representative images of P-Raf1-S259 staining. **(D)** Quantification of percent positive of total cells for P-Raf1-S259 and MFI per cell showing significantly more cells with phosphorylated Raf despite high ERK signaling. **(E)** ERK/Akt double negative and double positive percent of total cells showing loss of signaling insulation within Myr-Akt wells. (**F)** Representative image of stem cell (OLFM4, yellow) and TA cell (Ki-67, blue) markers in control and Myr-Akt cells. OLFM4+ cells lose proper stem cell niche patterning and instead exhibit a dysplastic phenotype. **(G)** Quantification showing extreme upregulation of OLFM4+ stem cells and TA cells in Myr-Akt wells. **(H)** Model depicting Myr-Akt causing constitutive hyperactivity of Akt and ERK pathways simultaneously leading to a hyperactive, dysplastic phenotype. Percent positive data shown is from analysis of 4 technical replicates with at least 150 total cells quantified per replicate. MFI quantifications are of hundreds of cells within one biological replicate, at least three total biological replicates were performed. Data are represented as mean ± SEM. All scale bars are 100μm, significance calculated with Welch's t-test, **** $P \leq 0.0001$

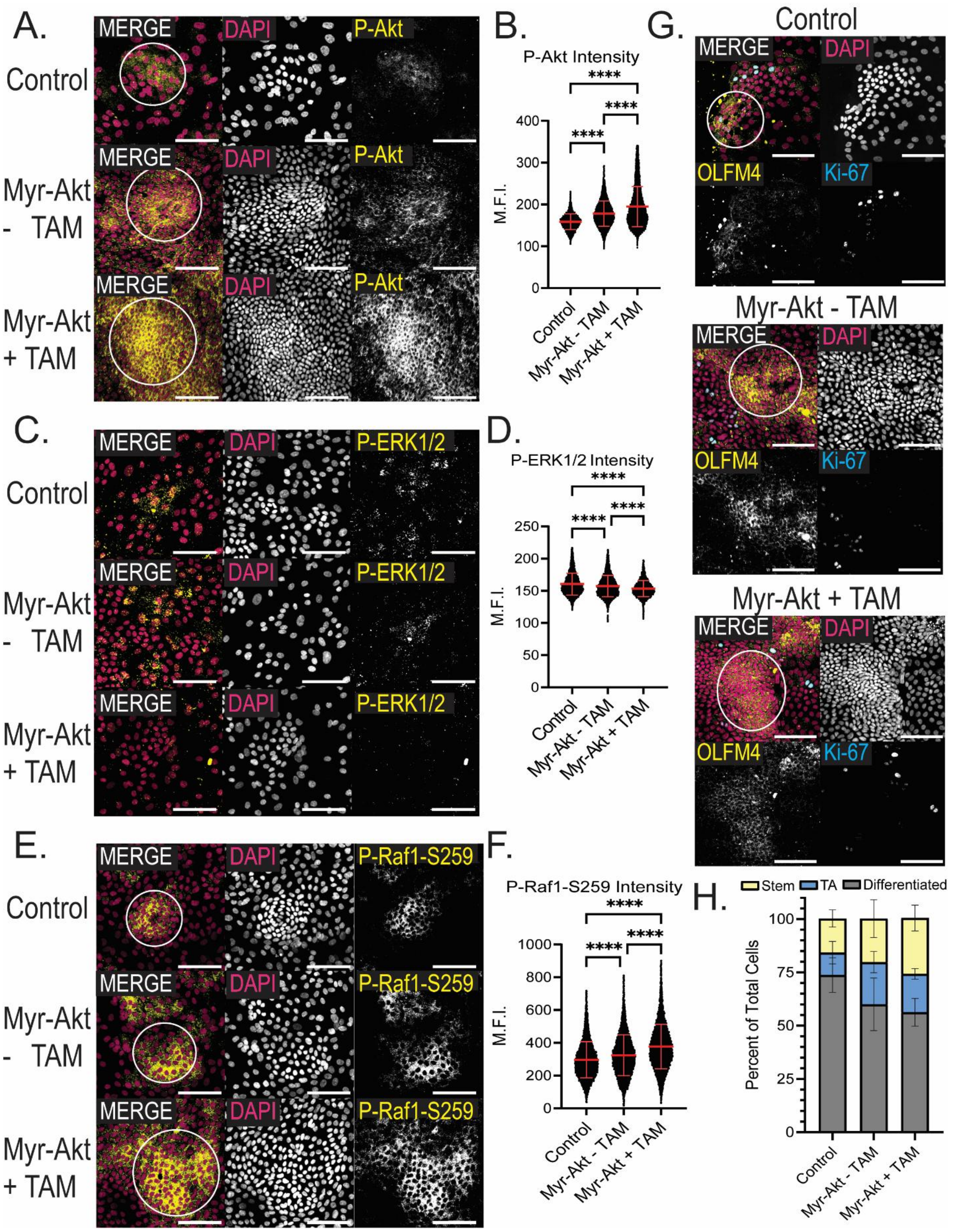
A.
MERGE
DAPI
P-Akt
Control
Myr-Akt - TAM
Myr-Akt + TAM
B.
P-Akt Intensity
M.F.I.
****
C.
P-ERK1/2
D.
P-ERK1/2 Intensity
E.
P-Raf1-S259
F.
P-Raf1-S259 Intensity
G.
OLFM4
Ki-67
H.
Stem
TA
Differentiated
Percent of Total Cells

**Figure Supplement 7**: **Temporally regulated AKT activation maintains signaling insulation**, related to Figure 5

**(A)** Representative image of control cells and cells infected with lentiviral Tamoxifen- inducible myristoylated Akt-473D construct causes hyperactivity of Akt in a dose-dependent manner. Construct still showed increased activity without induction, however addition of 1μM 4-Hydroxytamoxifen still induced further Akt activation. **(B)** Quantification of mean fluorescent intensity (MFI) of P-Akt in control, inducible Myr-Akt-473D without Tam, and Myr-Akt-473D with Tam treated wells. **(C)** Control and Myr-Akt representative images of P-Erk1/2 staining. **(D)** Quantification of P-ERK1/2 MFI showing a dose-dependent decrease in ERK signaling with induction of Myr-Akt-473D. **(E)** Control and Myr-Akt representative images of P-Raf1-S259 staining. **(F)** Quantification of P-Raf-S259 MFI showing a dose-dependent increase in Raf phosphorylation. **(G)** Representative image of stem cell (OLFM4, yellow) and TA cell (Ki-67, blue) markers in control and Myr-Akt-573 induced and non-induced cells, showing an increase in stem cell niches that maintain proper patterning. **(H)** Quantification showing a dose-dependent increase in stem and TA populations. Percent positive data shown is from analysis of 4 technical replicates with at least 150 total cells quantified per replicate. MFI quantifications are of hundreds of cells within one biological replicate, at least three total biological replicates were performed. Data are represented as mean ± SEM. All scale bars ate 100μm. Significance calculated with ANOVA test, **** $P \leq 0.0001$

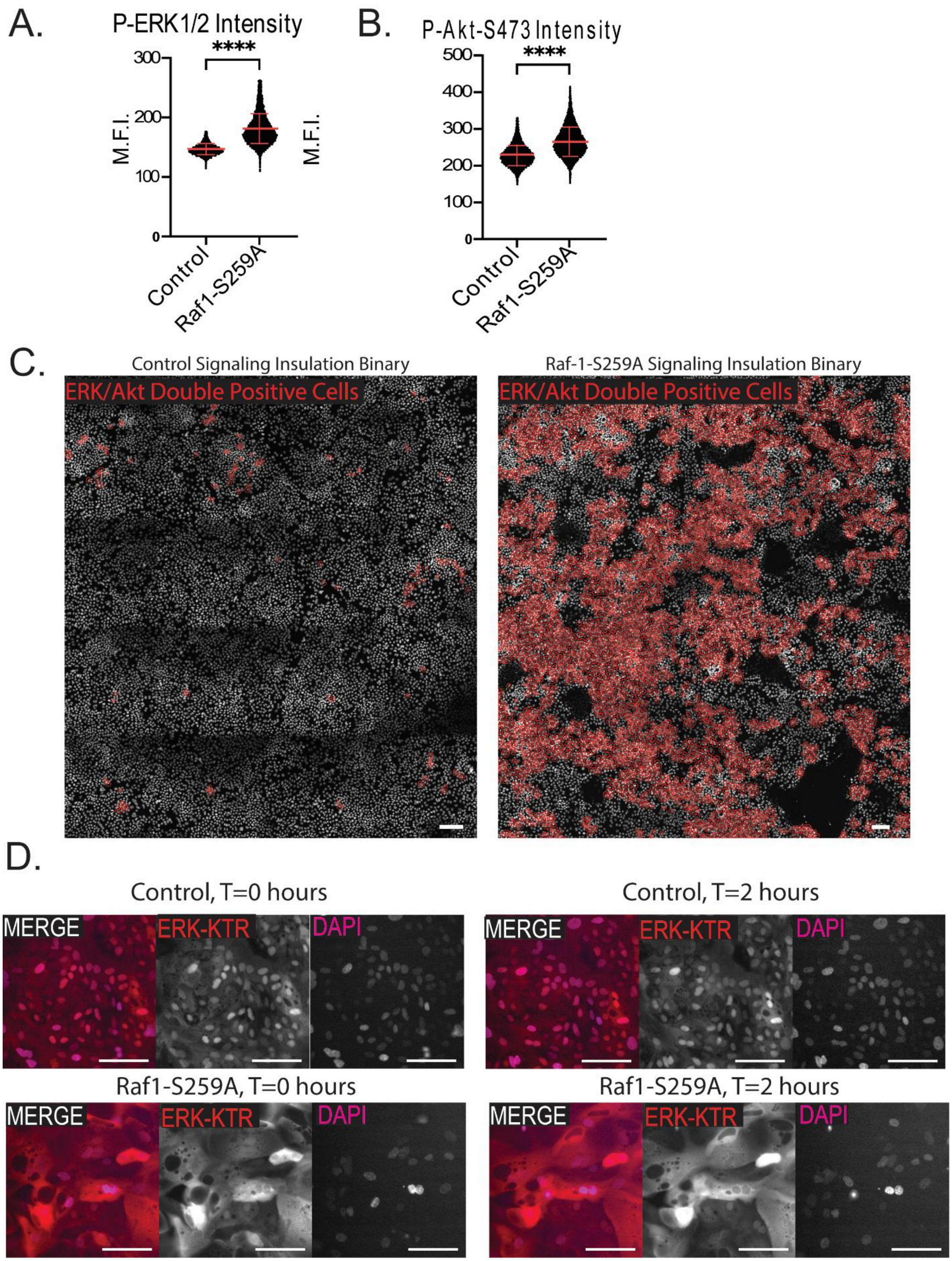
A.
P-ERK1/2 Intensity
****
M.F.I.
300
200
100
0
Control
Raf1-S259A
B.
P-Akt-S473 Intensity
****
M.F.I.
500
400
300
200
100
0
Control
Raf1-S259A
C.
Control Signaling Insulation Binary
ERK/Akt Double Positive Cells
Raf-1-S259A Signaling Insulation Binary
ERK/Akt Double Positive Cells
D.
Control, T=0 hours
MERGE
ERK-KTR
DAPI
Control, T=2 hours
MERGE
ERK-KTR
DAPI
Raf1-S259A, T=0 hours
MERGE
ERK-KTR
DAPI
Raf1-S259A, T=2 hours
MERGE
ERK-KTR
DAPI

**Figure Supplement 8**: **ERK Checkpoint is required for signaling insulation**, related to Figure 6

**(A)** Quantification P-ERK1/2 intensity in control and Raf1-S259A mutant wells, shown as the mean fluorescent intensity (MFI) of each cell in the monolayer. **(B)** Quantification of P-Akt-473 in control and Raf mutant wells, shown as the MFI of each cell in the monolayer. **(C)** Representative image of binary used for signaling insulation graphs, where red cells are ERK/Akt double positive and therefore uninsulated showing a drastic increase in uninsulated cells due to Raf1-S259A mutation. **(D)** Representative images of ERK-KTR organoids in control groups or Raf mutant cells 2 hours after start of movie. Raf mutant cells show a higher baseline activation and loss of on/off dynamics, staying constitutively activated. MFI quantifications are of hundreds of cells within one biological replicate, at least three total biological replicates were performed. Data are represented as mean ± SEM. All scale bars are 100μm, significance calculated with ANOVA test, **** P ≤ 0.0001.

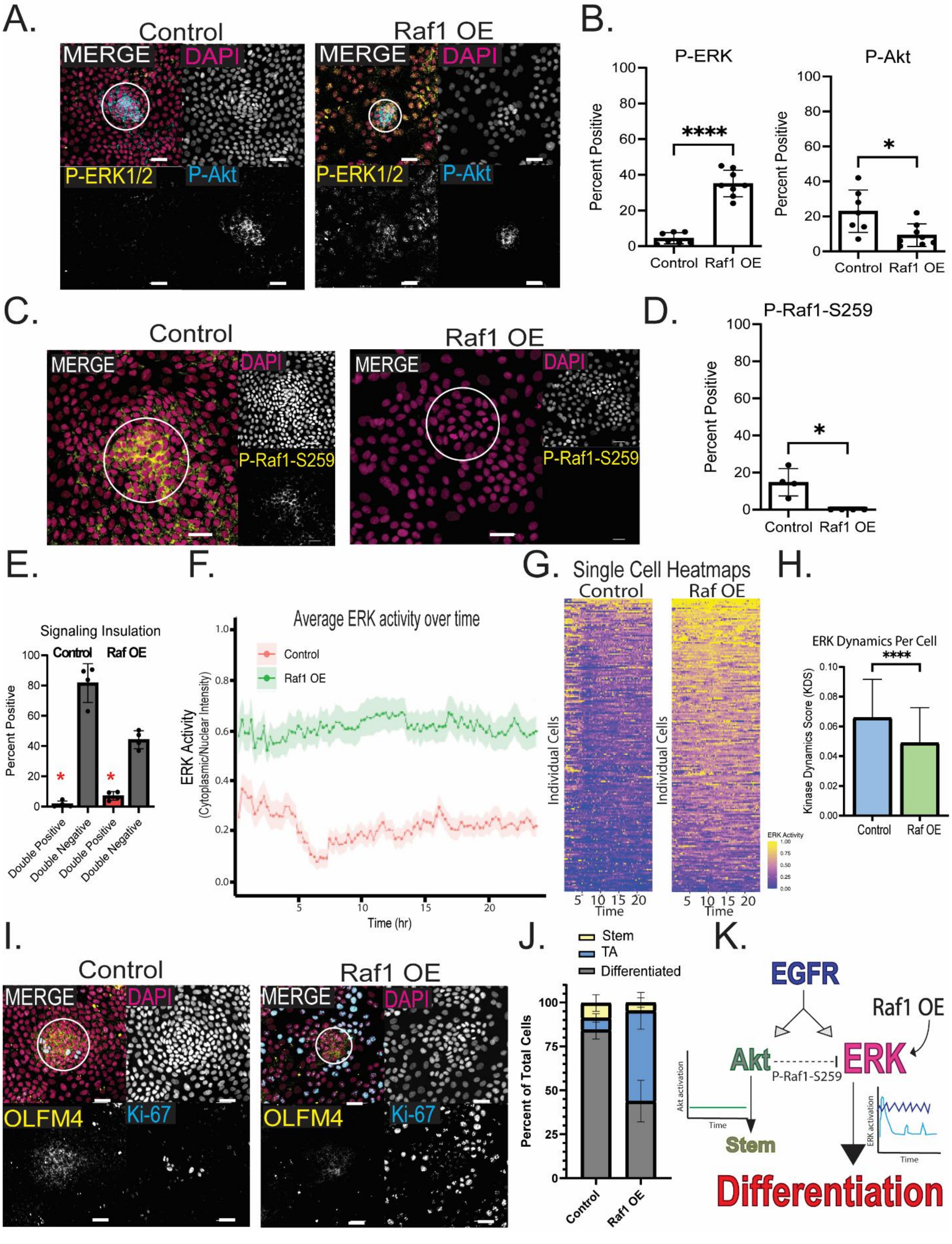
A.
Control
Raf1 OE
MERGE
DAPI
P-ERK1/2
P-Akt
B.
P-ERK
P-Akt
Percent Positive
****
*
Control
Raf1 OE
C.
Control
Raf1 OE
MERGE
DAPI
P-Raf1-S259
D.
P-Raf1-S259
Percent Positive
*
Control
Raf1 OE
E.
Signaling Insulation
Control
Raf OE
Percent Positive
Double Positive
Double Negative
Double Positive
Double Negative
F.
Average ERK activity over time
Control
Raf1 OE
ERK Activity
(Cytoplasmic/Nuclear Intensity)
Time (hr)
G.
Single Cell Heatmaps
Control
Raf OE
Individual Cells
Time
ERK Activity
H.
ERK Dynamics Per Cell
****
Kinase Dynamics Score (KDS)
Control
Raf OE
I.
Control
Raf1 OE
MERGE
DAPI
OLFM4
Ki-67
J.
Stem
TA
Differentiated
Percent of Total Cells
Control
Raf1 OE
K.
EGFR
Raf1 OE
Akt
ERK
P-Raf1-S259
Akt activation
Time
ERK activation
Time
Stem
Differentiation

**Figure Supplement 9: RAF-1 OE maintains signaling insulation and dynamics despite high kinase signaling load,** related to Figure 6

**(A)** Representative image of control cells and cells infected with lentiviral Raf1 overexpression construct showing global induction of ERK and reduction in Akt activity. Lentivirus was added to single cells during monolayer preparation due to difficulties growing the organoids in 3D. **(B)** Quantification of percent positive of total cells for P-ERK and P-Akt showing significant hyperactivity of ERK with lower activity of Akt. **(C)** Control and Raf OE representative images of P-Raf1-S259 staining. **(D)** Quantification of percent positive of total cells for P-Raf1-S259 showing a complete loss of Raf phosphorylation throughout the well. **(E)** Quantification of ERK/Akt double positive cells and double negative cells. Red asterisks indicate less than 5% of cells activate both pathways, signaling insulation which is maintained in the presence of PMA. **(F)** Quantification of ERK-KTR biosensor in organoids treated with PMA and control. Organoids were treated with 100nM PMA 1 hour after starting the movie. Graph shows average ERK activity by cytoplasmic/nuclear ration of KTR intensity with ribbon showing 95% CI. **(G)** Heatmaps showing ERK activity of all cells tracked, with each cell being a horizontal line on the graph. Yellow shows high ERK activity and blue being low activity. **(H** Kinase dynamics score (KDS) violin plot calculated as the standard deviation of ERK activity within each cell over the first 5 hours of the movie. **(I)** Representative image of stem cell (OLFM4, yellow) and TA cell (Ki-67, blue) markers in control and Raf1 overexpression cells. **(J)** Quantification showing extreme increase in TA cells with an almost total loss of stem cells. **(K)** Model depicting Raf OE causing a global increase in ERK activity that maintains dynamics while suppressing Akt signaling causing a loss of stemness and an increase in TA cell fates. Percent positive data shown is from analysis of 4 technical replicates with at least 150 total cells quantified per replicate. Data are represented as mean ± SEM. All scale bars are 100μm, significance calculated with Welch's t-test, * $P \leq 0.05$, **** $P \leq 0.0001$

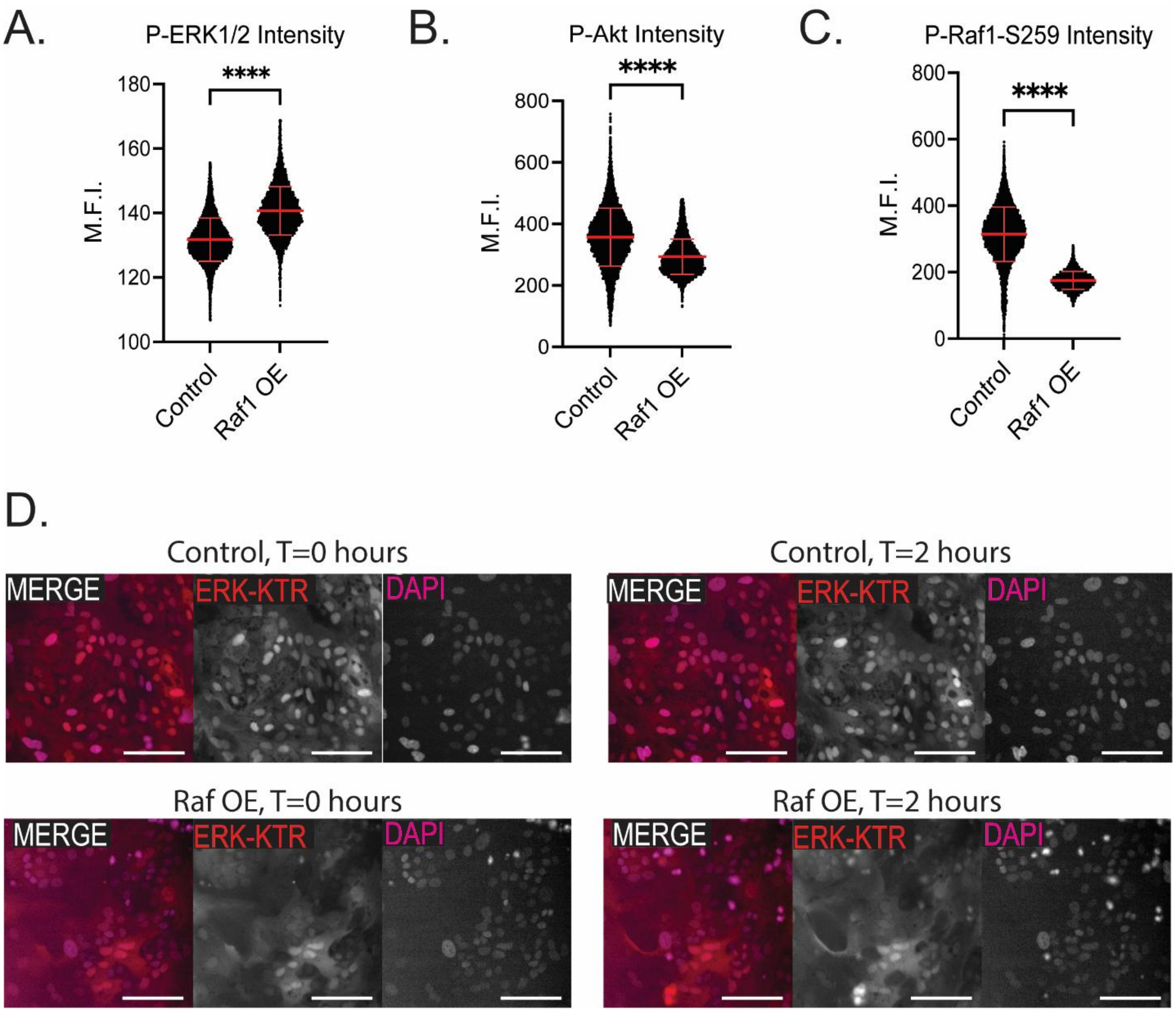


**Figure Supplement 10: Raf-1 overexpression maintains signaling insulation and dynamics**, related to Figure 6

**(A)** Quantification P-ERK1/2 intensity in control and Raf overexpression wells, shown as the mean fluorescent intensity (MFI) of each cell in the monolayer. **(B)** Quantification of P-Akt-473 in control and Raf overexpression wells, shown as the MFI of each cell in the monolayer. **(C) )** Quantification of P-Raf-S259 in control and Myr-Akt wells, shown as the MFI of each cell in the monolayer. **(D)** Representative images of ERK-KTR organoids in control groups or Raf overexpression cells 2 hours after start of movie. Raf overexpression cells show a higher baseline activation but still maintain on/off dynamics. MFI quantifications are of hundreds of cells within one biological replicate, at least three total biological replicates were performed. Data are represented as mean ± SEM. All scale bars are 100µm, significance calculated with ANOVA test, **** P ≤ 0.0001.

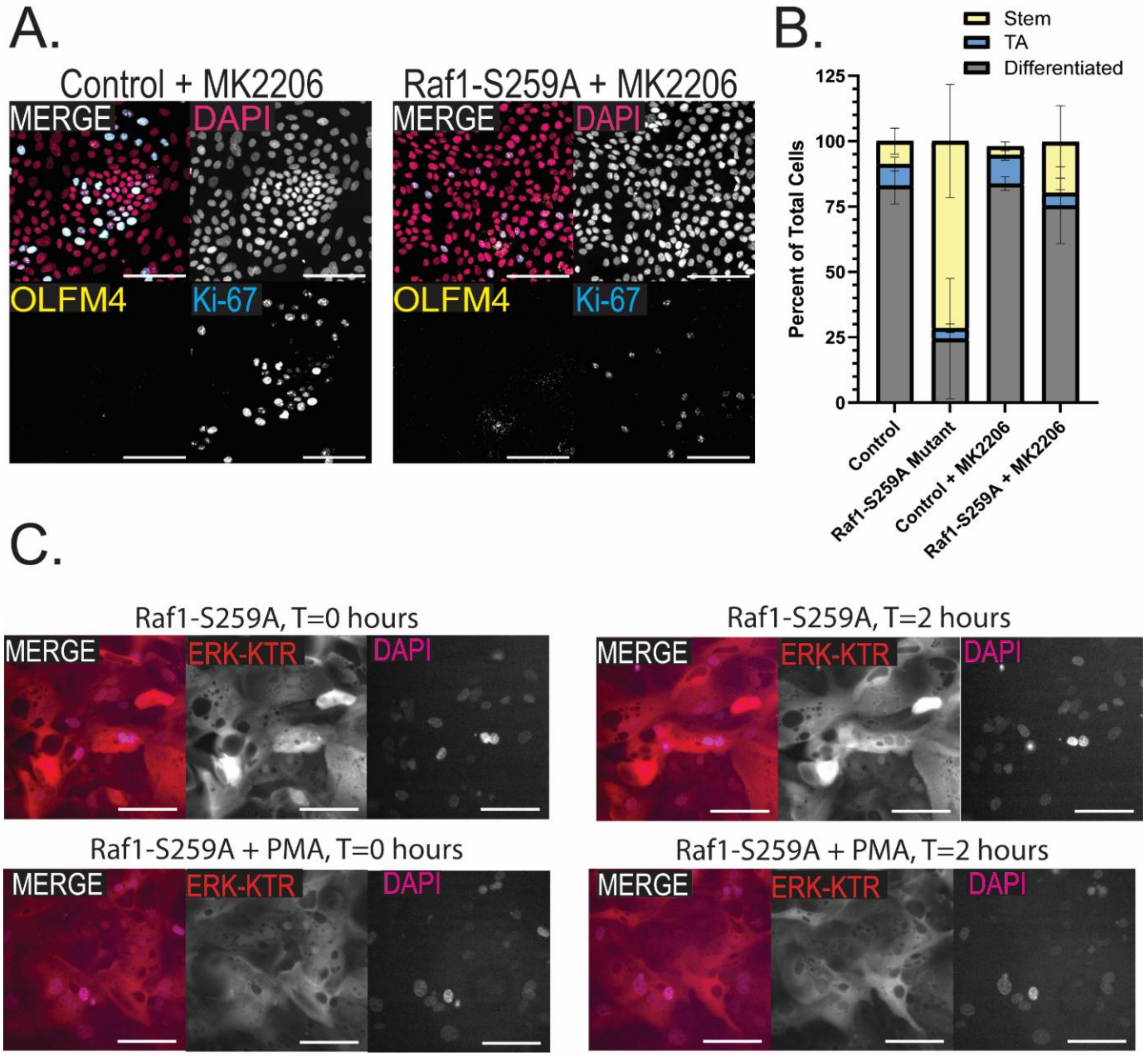


**Figure Supplement 11**: **AKT inhibition rescues neoplastic cell fate**, related to Figure 7

**(A)** Representative images of Control, Raf1-S259A phosphomutant, and both conditions treated with 1uM MK2206 for 72 hours. All wells were stained with OLFM4 (stem cell marker, yellow) and Ki-67 (TA cell marker, blue). **(B)** Quantification of all four conditions showing addition of MK2206 to Raf mutant cells can partially rescue the phenotype and cause global differentiation without increasing TA cell populations seen in control treated wells. **(C)** Representative images of ERK-KTR organoids in control groups or Raf mutant cells 2 hours after treatment with PMA. Raf mutant cells show a further increase in ERK activity, even with high baseline activity at start. All scale bars are 100 µm, data shown is from analysis of 4 technical replicates with at least 150 total cells quantified per replicate. Data are represented as mean ± SEM.
